# hiPSC-derived astrocytes from individuals with schizophrenia induce a dystrophic phenotype in microglial-like cells

**DOI:** 10.1101/2024.09.20.614161

**Authors:** Pablo L. Cardozo, Chia-Yi Lee, Juliana P. S. Lacerda, Júlia S. Fahel, Pablo Trindade, Gabriela Vitória, Leonardo Chicaybam, Rafaela C. Cordeiro, Isaque J. S. de Faria, Nathália C. Silva, Yaovi M. H. Todjro, Joana C. do P. Maciel, Martin H. Bonamino, Luciene B. Vieira, Breno F. Cruz, Rodrigo Nicolato, Kristen J. Brennand, Stevens K. Rehen, Fabíola M Ribeiro

**Affiliations:** Department of Biochemistry and Immunology, Institute of Biological Sciences (ICB), Universidade Federal de Minas Gerais (UFMG), Belo Horizonte, MG, Brazil; Department of Genetics, Yale School of Medicine, Yale University, New Haven, CT, United States of America; D’or Institute for Research and Education (IDOR), Rio de Janeiro, RJ, Brazil; Molecular Carcinogenesis Program, Research Coordination, Instituto Nacional do Câncer (INCA), Rio de Janeiro, RJ, Brazil; Vice-Presidency of Research and Biological Collections (VPPCB), Fundação Oswaldo Cruz (FIOCRUZ), Rio de Janeiro, RJ, Brazil; Department of Pharmacology, Institute of Biological Sciences (ICB), Universidade Federal de Minas Gerais (UFMG), Belo Horizonte, MG, Brazil; Department of Psychiatry, Faculty of Medicine, Universidade Federal de Minas Gerais (UFMG), Belo Horizonte, MG, Brazil; Department of Psychiatry, Yale School of Medicine, Yale University, New Haven, CT, United States of America; Department of Genetics, Institute of Biology, Universidade Federal do Rio de Janeiro (UFRJ), Rio de Janeiro, RJ, Brazil; Department of Clinical Analysis and Toxicology, Faculty of Pharmacy, Universidade Federal do Rio de Janeiro (UFRJ), Rio de Janeiro, RJ, Brazil

**Author notes:** These authors contributed equally to this work (Chia-Yi Lee, Juliana P. S. Lacerda).

**Keywords:** schizophrenia, astrocytes, microglia, dystrophic, CX3CL1, CX3CR1

## Abstract

Neuroinflammation, particularly astrocyte reactivity, is increasingly linked to schizophrenia (SCZ). Yet, the crosstalk between astrocytes and microglia in SCZ, especially under pro-inflammatory conditions, remains unclear. Here, we employed human induced-pluripotent stem cells to compare how astrocytes from five age-matched individuals with SCZ and five neurotypical controls, upon stimulation with TNF-α, affected microglial biology. TNF-ɑ stimulation of SCZ astrocytes, relative to their control counterparts, triggered increased mRNA expression of pro-inflammatory cytokines and CX3CL1. Interestingly, transcriptomic and gene set enrichment analyses revealed that reactive SCZ astrocytes promoted the downregulation of biological processes associated with immune cell proliferation and activation, phagocytosis and cell migration in induced microglial-like cells (iMGs). Under such conditions, iMGs assumed a dystrophic/senescent-like phenotype, which was associated with accelerated transcriptional aging. Functional validations showed that TNF-ɑ-stimulated SCZ astrocytes promoted reduced synaptoneurosomes phagocytosis by iMGs. Interestingly, only reactive control astrocytes were capable to induce significant microglial migration in a CX3CR1-dependent manner, despite the greater CX3CL1 secretion by reactive SCZ astrocytes compared to their stimulated control counterparts. This was likely due to SCZ astrocytes triggered reduction in CX3CR1 plasma membrane levels in iMGs. Altogether, these findings suggest that astrocytes contribute to SCZ pathology by altering normal microglial function and inducing a dystrophic phenotype.

**Main points:**

- SCZ astrocytes display an increased pro-inflammatory profile upon stimulation with TNF-α.
- Reactive SCZ astrocytes induce microglial-like cells to assume a dystrophic phenotype.
- Reactive SCZ astrocytes secreted factors impair microglial-like cells synaptoneurosome phagocytosis and limit their migration.

## INTRODUCTION

Schizophrenia (SCZ) is a neurodevelopmental disorder with an estimated lifetime prevalence ranging between 0.3-0.7% worldwide (Solmi et al., 2023). Symptoms usually appear between late adolescence and early adulthood (Hafner, Maurer, Loffler, & Riecher-Rossler, 1993) and can be divided into three broad categories: positive symptoms, such as hallucinations, psychosis and delusions; negative symptoms, e.g., depressive behavior, distraught thoughts and social withdrawn; and cognitive deficits, including working memory impairments, learning disabilities and attention deficits (Cannon, 2015; Perez & Lodge, 2014; Sakurai et al., 2015).

The causes of SCZ remain poorly understood, as both genetic and environmental risk factors interact to generate susceptible conditions towards the onset of this multifactorial disorder later in life (Hilker et al., 2018; Lipner, Murphy, & Ellman, 2019; Muller, Weidinger, Leitner, & Schwarz, 2015; Nimgaonkar, Prasad, Chowdari, Severance, & Yolken, 2017). Environmental risk factors, including childhood stress, obstetric complications and maternal infection during pregnancy (Lipner et al., 2019; Nimgaonkar et al., 2017), trigger inflammatory signaling cascades, which affect the developing brain at prenatal and early postnatal stages (Hilker et al., 2018; Muller et al., 2015). Indeed, genetic, transcriptional and serological evidence indicate increased inflammatory conditions in patients with SCZ, given the enhanced levels of multiple cytokines, such as IL-1β, IL-6, IL-8 and TNF-α (Frydecka et al., 2018; Muller et al., 2015; Rodrigues-Amorim et al., 2018). Moreover, genetic variants associated with SCZ show dynamic regulatory activity in neural cells in response to cytokine exposure (Retallick-Townsley et al., 2024).

Astrocytes provide metabolic support to neurons, facilitating the formation and modulation of synaptic connectivity, ionic buffering, while also uptaking and releasing neurotransmitters (Bernaus, Blanco, & Sevilla, 2020). Along with microglia, astrocytes perceive pro-inflammatory stimulation through a wide range of immune receptors, ultimately leading to their activation (Linnerbauer, Wheeler, & Quintana, 2020). Astrocyte activation follows enhanced expression of reactivity markers, such as GFAP and Vimentin, as well as augmented secretion of a wide range of cytokines and chemokines (e.g., TNF-α, IL-6, GM-CSF, TGF-β, CCL2) (Bernaus et al., 2020; Linnerbauer et al., 2020; Liu, Tang, & Feng, 2011). Secretion of these immune factors can, in turn, act upon microglial cells, either boosting or suppressing their activation, greatly impacting their effector capabilities, such as phagocytosis and cell migration (Bernaus et al., 2020; Linnerbauer et al., 2020; Liu et al., 2011). Astrocytes also modulate microglial function under physiological conditions; for instance, astrocyte-produced IL-33 stimulates synaptic pruning by microglia, which is critical for proper neuronal connectivity (Vainchtein et al., 2018).

Human genetic, post-mortem, and brain imaging analyses, together with mouse behavioral studies, link astrocytic dysfunction to SCZ (de Oliveira Figueiredo, Cali, Petrelli, & Bezzi, 2022; Dietz, Goldman, & Nedergaard, 2020). Astrocyte and interferon-response gene expression modules were upregulated in patients with SCZ, while the microglia module was downregulated (Gandal et al., 2018). Likewise, PET-scans reveal increased astrocyte reactivity in individuals with SCZ in the anterior cingulate cortex and left hippocampus (Kim et al., 2024). Finally, human induced pluripotent stem cells (hiPSCs)-derived astrocytes from individuals with SCZ have shown impaired astrocytic differentiation and maturation (Windrem et al., 2017), abnormal calcium signaling (Szabo et al., 2021), decreased glutamate uptake (Szabo et al., 2021), and enhanced pro-inflammatory profile (Koskuvi et al., 2022; Trindade et al., 2023).

Nonetheless, the crosstalk between astrocytes and microglia in SCZ, especially under pro-inflammatory conditions, remains unclear. Here we evaluated how reactive SCZ hiPSC-derived astrocytes affected the microglial phenotype, both at the transcriptional and functional levels. We further assessed the contribution of CX3CL1/CX3CR1 axis in this phenomenon and aimed at identifying putative transcriptional factors driving the observed alterations.

## METHODS

### Ethics Statement

All experiments carried out throughout this work were approved by UFMG’s Ethics Committee (COEP-UFMG #90424518.3.1001.5149) and were performed according to the Helsinki Declaration and the Brazilian National Health Council Resolution 466/12.

### Resources Table

A complete list of all materials and reagents used in this study can be found in **Table S1**.

### Candidate Genes Screening

The list of transcripts or proteins identified in two transcriptomic (Akkouh et al., 2022; Szabo et al., 2021) and one proteomic (Trindade et al., 2023) studies comparing gene expression between hiPSC-derived astrocytes sourced from individuals with schizophrenia and controls under basal conditions were obtained from each study supplementary data. These lists were inputted into *Metascape* (Zhou et al., 2019) and up to 40 GO Biological Processes (GO BP) terms retrieved per study (p < 0.01). Overlapping GO BP terms were identified, and genes associated with the chosen GO BP term were obtained from each study to shortlist potential candidates. CX3CL1 associated GO BP were retrieved using the *GOxploreR* package (v. 1.2.7) (Manjang, Tripathi, Yli-Harja, Dehmer, & Emmert-Streib, 2020) and their semantic similarity matrix used for UMAP analysis in the *rrvgo* package (v. 1.12.2) (Sayols, 2023) in R (4.3.1). Synaptic pruning network figure was created by merging GO:0098883 (“synaptic pruning”) and “Complement Signaling” (downloaded from *SIGNOR 3.0*) and editing the resulting network in *Cytoscape* (v. 3.10.1) (Shannon et al., 2003) for schematic visualization.

### hiPSC-derived neural stem cells (NSCs)

Neural stem cells (NSCs) were differentiated from hiPSCs from five neurotypical and five individuals with schizophrenia (**Table S2**) as previously described (de Lima et al., 2023; Trindade et al., 2020; Yan et al., 2013). Briefly, 80-90% confluent hiPSCs were dissociated in single cells using Accutase solution and plated at 3 x 10^4^ cells/cm^2^ in Geltrex-coated tissue culture (TC)-treated 6-well plates in StemFlex Medium, supplemented with 1% antimycotic-antibiotics solution (anti-anti) and 10 µM Rho-kinase inhibitor Y-27632 (ROCKi). In the first day of differentiation (day 0), media was switched to PSC Neural Induction Medium (Neurobasal, 2% Neural Induction Supplement a 1% penicillin-streptomycin (P/S) solution) and changed every other day. On the 7th day, Neural Stem Cells (NSCs) were dissociated with Accutase and plated at 1 x 10^5^ cells/cm^2^ in Geltrex-coated TC 60-mm petri dishes in NSC Expansion Medium (50% Advanced DMEM/F-12, 50% Neurobasal, 2% Neural Induction Supplement and 1% P/S) with 10 µM ROCKi. Media changes were performed three times a week. Cells were cultured in standard conditions (5% CO2 atmosphere and 37 °C in a humidified incubator). hiPSCs identity was confirmed by positive immunostaining to OCT4, SSEA-1, TRA-1-60 and TRA-1-81 (**Figures S1A-D**). NSCs identity was confirmed by positive immunostaining to Nestin, PAX6, SOX1 and SOX9 (**Figures S1E-H**).

### hiPSC-derived astrocytes differentiation

Astrocytes differentiation was performed as described elsewhere (de Lima et al., 2023; Trindade et al., 2020). Once hiPSC-derived neural stem cells (NSCs) reached 90% confluency, cells were dissociated with Accutase and seeded at 5 x 10^4^ cells/cm^2^ in Geltrex-coated TC 25 cm^2^ flasks in Astrocytes Differentiation Medium (DMEM/F-12, 1x N-2 supplement, 1% Heat-inactivated Fetal Bovine Serum (FBS) and 1% antimycotic-antibiotics solution (anti-anti)). Media was replenished every other day for 21 days. Whenever differentiating astrocytes reached 90% confluency, cells were expanded to larger Geltrex-coated TC flasks at 1:3 ratio as described. By the end of this differentiation period (day 21), media was switched to Astrocytes Maturation Medium (DMEM/F-12, 10% FBS and 1% anti-anti) to mature astrocytes for 5 weeks. Whenever astrocytes reached 90% confluency, they were split using Trypsin/EDTA 0.125% solution (diluted in PBS 1x^-/-^) and expanded to 175 cm^2^ TC flasks without Geltrex coating at 1:2 ratio to negatively select undifferentiated non-adhering cells. Cells were cultured in standard conditions. Astrocytes’ identity was confirmed by positive immunostaining to EAAT1, Vimentin, GFAP and Connexin 43 **(Figures S2A-B).**

### Astrocytes Stimulation

After at least 5 weeks of maturation, astrocytes were detached using Trypsin/EDTA 0.125% solution, pelleted down by centrifugation (800 g for 5 min), resuspended in Astrocytes Maturation Medium and plated at 1.25 x 10^4^ cell/cm^2^ density in either TC 6-well plates or 25 cm^2^ flasks. After 5 days in culture, cells were washed three times with PBS 1^-/-^ and serum-starved for 24 h in DMEM/F-12 supplemented with 1% anti-anti. Next, astrocytes were stimulated for 24 h with 10 ng/mL recombinant human TNF-α or vehicle (0.1% BSA solution). Once stimulation finished, astrocytes conditioned media (A_CM_) was collected, clarified by centrifugation (3000 g, 4 °C for 10 min), flash frozen and stored at −80 °C until further use. In addition, astrocytes monolayers were harvested either in TRIzol reagent according to manufacturer instructions or in RIPA lysis buffer containing protease and phosphatase inhibitors for 1 h on ice and stored at −80 °C.

### Total RNA extraction and RT-qPCR

Total RNA was isolated using the TRIzol reagent as per manufacturer instructions and resuspended in 12 µL DEPC-treated nuclease-free water. RNA concentration and quality (260/230 and 260/280 ratios) were analyzed by spectrophotometry in the Multiskan GO (Thermo Scientific). 1 µg total RNA was reverse transcribed in a reaction mixture consisting of 15 ng/µL Random Primers, 50 mM Tris-HCl, 75 mM KCl, 3 mM MgCl_2_, 625 µM dNTPs, 10 µM DTT and M-MLV reverse transcriptase. Reverse transcription conditions were as follows: (Random primers annealing) 70 °C for 10 min and 4 °C for 10 min; (cDNA extension) 42 °C for 60 min and 70 °C for 15 min. The resulting cDNA was diluted 1:10 in nuclease-free water and subjected to qPCR using Power SYBR Green PCR Master Mix and 200 nM of forward and reverse primers mix (sequences available in **Table S3**) in the QuantStudio 7 System (Applied Biosystems). Thermal cycling conditions were: 50 °C for 2 min, 95 °C for 10 min; 40 cycles of 95°C for 15 s and 60 °C for 1 min; followed by melting curve analysis. All primers pairs had 90-110% efficiency. Relative gene expression of target genes was normalized by the average of housekeeping genes (RPLP0 and IPO8) and calculated by the 2-^ΔCt^ method.

### Donor selection and peripheral-blood mononuclear cells (PBMCs) isolation

A male control individual was chosen (**Table S2**), after evaluation by a trained psychiatrist following the M.I.N.I. 7.0.0 (Sheehan et al., 1998) and fitting the following inclusion and exclusion criteria: less than 50 years of age; at least high school education; no previous history of neurological or neuropsychiatric conditions; absence of 1^st^ degree relatives diagnosed with psychiatric disorders; absence of 2^nd^ degree relatives diagnosed with schizophrenia, bipolar disorder or autism spectrum disorder; non-cannabis user; non-smoker; capability to freely and voluntarily give informed consent to participate in this study. The study design was fully and carefully explained to the selected participant, who had the chance to ask questions afterwards to clarify any concern regarding his study participation. Next, both the participant and the corresponding author signed two copies of the Informed Consent Form before proceeding with the his blood harvest. Whole blood was drawn from this individual, diluted 1:1 with PBS 1x^-/-^, carefully layered onto Histopaque 1077 density gradient and centrifuged at 900 g for 25 min (with acceleration and braking settings turned off). After blood fractionation, the PBMCs layer was collected using a sterile plastic Pasteur pipette, resuspended in PBS 1x^-/-^ and centrifuged at 400 g for 8 min. The supernatant was removed, cells resuspended once again in PBS 1x^-/-^ and pelleted by centrifugation at 400 g for 8 min. PBMCs were resuspended in FBS, counted in a hemocytometer and cryopreserved in FBS + 10% DMSO for later use.

### Induced microglial-like cells (iMGs) differentiation

iMGs differentiation was carried out as described elsewhere (Ohgidani et al., 2014; Sellgren et al., 2019; Sellgren et al., 2017). PBMCs were plated in either 13-mm acid-etched glass coverslips or TC-treated plasticware, both previously coated overnight with Geltrex, and cultured in RPMI 1640 medium, supplemented with 10% FBS and 1% penicillin-streptomycin (P/S) solution. Cell density was adjusted on an assay-dependent basis. In the following day (day 0), media was completely switched to iMG differentiation media (MDM), comprised of RPMI 1640, 1% Glutamax, 100 ng/mL IL-34, 10 ng/mL GM-CSF and 1% P/S. On the 6^th^ day of differentiation, half of the media was changed and replenished by fresh MDM. On day 9, the media was thoroughly removed, cells washed twice with pre-warmed RPMI 1640 medium to remove cell debris and fresh MDM added. iMGs were used for subsequent experiments between days 10-12 of differentiation. iMGs were kept in standard culture conditions. Microglial-like cells identity was confirmed by their ramified morphology and positive immunostaining to CX3CR1, Iba1 and Pu.1. Bright-field images were captured using the FLoid Cell Imaging Station (Thermo Fisher).

### hiPSC-derived neuronal differentiation

CF1 and CF2 NSCs were differentiated to neurons, according to Espuny-Camacho et al. (2013) with modifications. 2.5 x 10^4^ or 5 x 10^4^ NSCs/cm^2^ were seed on 13-mm acid-etched sterile glass coverslips or TC-treated 100-mm petri dishes, respectively, coated with 0.1% poly-ethylenimine (glass surface) or 0.001% poly-ornithine (plastic surface) solution and 2% Geltrex, in Default Defined Medium (DDM), consisting of DMEM/F-12, 2% B-27 supplement, 1% N-2 supplement, 1% Glutamax, 1% MEM non-essential amino acids solution, 1 mM Sodium Pyruvate, 0.1 mM 2-mercaptoethanol. 0.5 mg/mL BSA, 200 ng/mL L-ascorbic acid, 1 µg/mL laminin and 1% P/S, supplemented with 10 µM ROCKi. After two days in culture, the media was thoroughly removed and replenished by fresh DDM without ROCKi. Three days later (day 0), half of the media was replaced by Neurobasal/B-27 medium (Neurobasal, 2% B-27 supplement, 1% Glutamax, 200 ng/mL L-ascorbic acid, 1 µg/mL laminin and 1% P/S). Half of the media was changed every 3-4 days by an isovolumetric mixture of DDM and Neurobasal/B-27 media until day 60 of differentiation. At the end of differentiation, neuronal cultures were either stained to β-tubulin III, MAP2, S100β, MBP, Synaptotagmin 2 (SYT 2) and PSD-95 or used for synaptoneurosomes isolation. Cells were kept in standard culture conditions.

### Synaptoneurosomes isolation

Synaptoneurosomes isolation was performed as described by Sellgren et al. (2019) with minor modifications. After 60 days of differentiation, media was removed from neuronal cultures and replaced by non-stimulated A_CM_ from their isogenic counterparts 24 h before cell harvest. Next, media was thoroughly removed, and cells washed twice with ice-cold PBS 1x. 1 mL per 100-mm TC dish of ice-cold Synaptoneurosomes Isolation Buffer (SIB; 10 mM HEPES, 1 mM EDTA, 2 mM EGTA, 0.5 mM DTT and protease inhibitors; pH 7.0) (Villasana, Klann, & Tejada-Simon, 2006) was added and neurons gently scrapped. This cell suspension was transferred to a sterile 1.5 mL conical tube and centrifuged at 1,200 g for 10 min at 4°C. The supernatant (homogenate) was transferred to a new sterile 1.5 mL conical tube and centrifuged at 15,000 g for 20 min at 4 °C. Finally, the supernatant (cytosolic fraction) was removed, pelleted synaptoneurosomes resuspended in SIB with 5% DMSO and frozen at −80 °C until further use. Aliquots of each fraction were collected at each step for further characterization by Western Blot and stored at −80 °C. Freshly isolated synaptoneurosomes were also plated overnight on 13-mm acid-etched glass coverslips coated with 50 µg/mL poly-D-lysine solution and left to attach overnight under standard culture conditions for further immunostaining to Syntaxin 1 and Homer. Shortly before the phagocytosis assay, synaptoneurosomes were homogenized for up to 50 times in ice-cold PBS 1x using a sterile micro-capillary pipet tip, their concentration adjusted to 0.25 µg/µL and labelled with 2 µM Vybrant-CM-DiI, as per manufacturer instructions.

### Synaptoneurosomes phagocytosis assay

2.8 x 10^5^ PBMCs/cm^2^ per well were seeded in 24-well plates and differentiated as already described. On day 12, iMGs were incubated for 1 h with 2 ng/mL recombinant CX3CL1 or vehicle (0.1% BSA solution). Alternatively, iMGs were pre-treated with 5 µM AZD8797 or vehicle (DMSO) for 1 h; then, half of the media was removed and replenished by an equal volume of pooled A_CM_ and cells incubated for another hour (AZD8797 was also added again to ensure its concentration was kept at 5 µM throughout the whole experiment). Next, 1.875 µg of CM-DiI-labelled synaptoneurosomes were added to each well, plates gently shaken for thorough homogenization and incubated for 24 h in standard culture conditions in the Cytation 5 Cell Imaging Multi-mode Reader (BioTek; CAPI - ICB, UFMG, Brazil). Four equally spaced image sets (bright field and RFP channels) per well were captured every two hours. Experiments were conducted twice in duplicates to ensure reproducibility. After 24 h, images were retrieved and analyzed using the combination of the following open-source softwares: (1) *FIJI/ImageJ* (v. 1.54f) (Schindelin et al., 2012) for brightness/contrast adjustments and denoising; (2) *fastER* (v. 1.3.5) (Hilsenbeck et al., 2017) for cell segmentation (bright field channel); (3) *CellProfiler* (v. 4.2.1) (Stirling et al., 2021) for single-cell phagocytic index quantification. Phagocytic index was calculated as the ratio of red object area (μm^2^) inside a given cell by its total cell area (μm^2^). Orthogonal projection (z-step: 2 µm; number of steps: 20) was generated using *FIJI/ImageJ* from vehicle-treated (0.1% BSA) fixed samples (24 h time-point) on glass coverslips immunostained for Alexa 633 Phalloidin and Hoechst 33342 as described.

### Western Blot

Astrocytes’ lysates (50 µg) and synaptoneurosomes fractions (25 µg) protein concentration was quantified by the Bradford method. Samples were diluted in Laemmli Sample Buffer, heated at 95 °C for 5 min and later subjected to electrophoretic separation in 10% SDS-PAGE. Proteins were transferred to 0.45 µm nitrocellulose membranes, blocked with 5% BSA solution diluted in TBS with 0.1% Tween-20 (TBS-T) for 1 h at room temperature, followed by overnight incubation at 4 °C with primary antibodies diluted in 3% BSA solution in TBS-T: anti-CX3CL1 (1:500), anti-β-actin (1:5000), anti-Syntaxin 1 (1:200), anti-Homer (1:500) or anti-Vinculin (1:10000). In the following day, primary antibodies were removed, membranes washed three times with TBS-T for 5 min and incubated for 1 h at room temperature with secondary antibodies diluted in 3% free-fat milk solution in TBS-T: HRP-conjugated anti-mouse IgG (1:2500), HRP-conjugated anti-rabbit IgG (1:2500) and HRP-conjugated anti-goat IgG (1:2500). Next, secondary antibodies were removed, membranes washed with TBS-T as described, incubated for 5 min with the detection reagent for chemiluminescence reaction and images acquired using the ImageQuant LAS 4000 platform. When appropriate, densitometric analysis was carried out using *FIJI/ImageJ* and expressed as ratio of CX3CL1/β-actin protein levels relative to non-stimulated HCT astrocytes.

### Immunofluorescence staining

Cells and synaptoneurosomes were fixed for 15 min in 4% PFA solution (in PBS 1x). Next, samples were permeabilized for 10 min in 0.3% Triton X-100 diluted in PBS 1x (PBS-T) and blocked in 2% BSA solution (in PBS-T) for 1 h. Afterwards, cells were incubated overnight at 4 °C with primary antibodies diluted in blocking solution: anti-OCT4 (1:200), anti-SSEA-4 (1:100), anti-TRA-1-60 (1:100), anti-TRA-1-81 (1:100), anti-Nestin (1:400), anti-PAX6 (1:100), anti-SOX1 (1:200), anti-SOX9 (1:100), anti-GFAP (1:200), anti-Vimentin (1:500), anti-EAAT1 (1:100), anti-Connexin 43 (1:100), anti-MAP2 (1:500), anti-β-tubulin III (1:500), anti-S100β (1:200), anti-MBP (1:500), anti-Synaptotagmin 2 (1:100), anti-PSD95 (1:500), anti-Syntaxin 1 (1:100), anti-Homer (1:200), anti-CX3CR1 (1:500), anti-Iba1 (1:750) and/or anti-Pu.1 (1:200). In the following day, primary antibody solution was removed, samples washed three times for 5 min with PBS and incubated for 1 h, protected from light and at room temperature with the following secondary antibodies and dyes diluted in blocking solution: Alexa Fluor 488 anti-Mouse IgG (1:400), Alexa Fluor 546 anti-Rabbit IgG (1:300), Alexa Fluor 546 anti-Rat IgG (1:300), Alexa Fluor 555 anti-Goat IgG (1:500), Alexa Fluor 594 anti-Mouse IgG (1:400), Alexa Fluor 633 anti-Mouse IgG (1:500), Alexa Fluor 633 anti-Rabbit IgG (1:500), Alexa Fluor 633 Phalloidin (1:1000) and/or Hoechst 33342 (1:500). After secondary antibody incubation, samples were washed as described and mounted on clean glass slides using the Hydromount mounting medium, overnight and protected from light. Coverslips were sealed with nail polish in the following day. Images were acquired via confocal laser microscopy using the Nikon A1 microscope (Centre for Gastrointestinal Biology, UFMG, Brazil) or via wide-field microscopy using the Cytation 1 Cell Imaging Multimode Reader (Biotek; D’or Institute, Brazil). Images were analyzed on *FIJI/ImageJ*.

### Enzyme-linked Immunosorbent Assay (ELISA)

Soluble CX3CL1 was quantified in A_CM_ using the Human CX3CL1/Fractalkine DuoSet ELISA kit as per manufacturer instructions. Astrocytes monolayers of all cell lines were detached using Trypsin/EDTA 0.125% solution and counted in a hemocytometer immediately after A_CM_ harvest to normalize data by cell count.

### cDNA library preparation and sequencing

4.0 x 10^5^ PBMCs/cm^2^ per well were seeded in Geltrex-coated TC 12-well plates and differentiated as described. After 12 days of differentiation, half of the media was discarded and replaced by an equal volume of pooled A_CM_. 24 h later, media was completely removed, and cells harvested in TRIzol reagent. Total RNA extraction was performed as recommended by the manufacturer and RNA quality analyzed in the 2100 Bioanalyzer platform (Agilent). 200 ng of total RNA (RIN > 7) was subjected to rRNA depletion and cDNA library preparation using the NEBNext Ultra II Directional RNA Library Prep Kit, according to manufacturer instructions. cDNA library quality assessment and paired-end sequencing (2 x 100 pb) using the NextSeq2000 platform (Illumina) was performed by the LaCTAD (UNICAMP, Brazil). Samples had an average sequencing coverage of 30 million reads/library.

### Differential gene expression analysis

RNA-seq reads quality was assessed by *FastQC* (Andrews, 2010). Adaptor sequences were trimmed out and reads with less than 50 bp or Phred < 20 were removed using *Trimmomatic* (v. 0.40) (Bolger, Lohse, & Usadel, 2014). The *STAR* aligner (v. 2.7.11b) (Dobin et al., 2013) was used for mapping reads to the human reference genome (GRCh38), and *featureCounts* (v. 2.0.6) (Liao, Smyth, & Shi, 2014) was employed to count mapped reads. Features with fewer than 10 counts were excluded from further analysis. Batch effects were removed using the *RUVseq* package (residuals method; v. 1.34.0) (Risso, Ngai, Speed, & Dudoit, 2014). *edgeR* (v. 3.42.4) (Robinson, McCarthy, & Smyth, 2010) was employed for differential gene expression analysis (adjusted p-value < 0.05; |log_2_ FC| ≥ 0.5). Principal component analysis was performed using the *PCAtools* package (v. 2.12.0) (Kevin Blighe, 2024).

### Gene set enrichment analysis (GSEA)

For GSEA, lists of ranked genes were generated using the Signal-to-Noise ratio metric from the batch-corrected gene count matrix. Next, these ranked lists were subjected to GSEA using the *clusterProfiler* package (v. 4.8.3) (Wu et al., 2021) to identify enriched GO Biological Processes and KEGG Pathways. The GSEA function was also used to identify microglial phenotypic signatures against the Prater et al. (2023) single-nuclei RNA-seq dataset. This dataset consists of human dorsolateral prefrontal cortex microglia isolated from *post-mortem* brain tissue of individuals with Alzheimer’s Disease; signature analysis was restricted to individuals bearing the APOE3/APOE3 genotype to prevent potential biases associated with the APOE4 allele. The GSEA function was also used to evaluate cellular senescence induction or inhibition using the CellAge database (Avelar et al., 2020). Adjusted p-value threshold was set at 0.1.

### Weighted Gene Co-expression Network Analysis (WGCNA)

For WGCNA, the batch-corrected gene count matrix was used. Genes with fewer than 10 counts were excluded, and the top 10,000 genes with the highest variance were selected for network construction. A soft thresholding power of 11 was applied using the signed-hybrid network type. The minimum module size was set to 30, and modules were merged if their Spearman correlation coefficient exceeded 0.8. Correlation p-values were adjusted using the Benjamini and Hochberg method (adjusted p-value < 0.05). The *bioNERO* package (v. 1.8.7) (Almeida-Silva & Venancio, 2022) was used alongside *WGCNA* (v. 1.72-5) (Langfelder & Horvath, 2008) for analysis and visualization. GO term enrichment analysis was performed using the *biomaRt* package (v. 2.56.1) (Durinck et al., 2005).

### Master Regulator Analysis (MRA)

MRA analysis was carried out based on the pipeline published by Leng et al. (2022) with adaptations, using the *RTN* package (v. 2.24.0) (Fletcher et al., 2013). Firstly, for co-expression network reconstruction, 871 RNA-seq datasets were downloaded from ARCHS4 (Lachmann et al., 2018), whose sample descriptors were “Microglia” and “Human”. Batch effects across datasets were removed using the *Combat-seq* function of the *sva* package (v. 3.48.0) (Jeffrey T. Leek, 2024), gene counts log transformed using the *vst* function of the *DESeq2* package (v. 1.40.2) (Love, Huber, & Anders, 2014) and gene-level count matrix subset to contain only the 16,954 genes identified in the RNA-seq dataset of the present study. The list of human transcription factors published by Lambert et al. (2018) was input as regulatory elements for network reconstruction. After transcriptional regulatory network reconstruction finished, master regulators and their activity status for each experimental condition were calculated as described in the *RTN* package vignette (adjusted p-value < 0.1). Associations between activated or repressed regulons and WGCNA modules were done using the *tni.overlap.genesets()* function.

### Transcriptional age calculation

Transcriptional age was calculated from the batch-corrected gene count matrix by the *RNAAgeCalc* package (v. 1.16) (Ren & Kuan, 2020), using the “GTExAge” signature and “brain” as reference tissue.

### iMGs migration assay

3.8 x 10^5^ PBMCs/cm^2^ were plated in Geltrex-coated 60-mm TC petri dishes and differentiated as already described. After 9 days of differentiation, iMGs were pre-treated with 10 µM Rho-kinase inhibitor Y-27632 for 1 h, followed by incubation with hypertonic citrate saline solution (135 mM KCl and 15 mM sodium citrate diluted in PBS 1x) for 5 min at 37 °C to gently loosen up attached cells. Next, iMGs were gently scraped using a cell scraper, collected into a 15 mL conical tube (pre-coated with sterile 5% BSA solution, overnight at 4°C) and centrifuged at 400 g for 8 min. Supernatant was discarded, iMGs resuspended in MDM and 6.0 x 10^4^ cells/transwell plated into the upper compartment of each Geltrex-coated transwell. 24 h after plating, iMGs were pre-treated with AZD8797 (5 µM) or vehicle (DMSO) for 1 h, pooled A_CM_ was added in the lower compartment and cells were incubated overnight (∼ 16 h) in standard culture conditions. In the next day, iMGs migration was stopped by fixing these cells for 15 min with 4% PFA. Transwells were washed three times with PBS 1x, non-migrated cells (in the upper compartment) gently scraped off using a cotton swab and migrated cells (in the lower compartment) stained with Hoechst 33342 (1:500; diluted in PBS 1x) for 10 min and protected from light. Samples were washed as described and subsequently imaged in the EVOS FL Color microscope (Thermo Fisher). 4-5 image fields/transwell were randomly acquired and migrated cells quantified by a trained researcher, blind to the experimental conditions, using the *CellCounter* plugin on *FIJI/ImageJ*. Migrated cell count was expressed as percentage relative to vehicle-treated iMGs exposed to A_CM_ HCT (N.S.). This experiment was executed three times independently.

### Cell-based ELISA

5.26 x 10^5^ PBMCs/cm^2^ per well were plated in Geltrex-coated TC 96-well plates and differentiated to iMGs as described. Once differentiation was completed, half of the media was removed, replenished by an equal volume of pooled A_CM_ and incubated for 24 h in standard culture conditions. Afterwards, media was carefully removed, and cells fixed with 4% PFA solution (in PBS 1x) for 15 min. Then, cell-based ELISA was carried out based on Yang, Zhou, Roberts, Nie, and Tang (2021) with modifications. Samples were washed three times with PBS 1x for 5 min under gentle agitation to remove excessive fixative solution. Next, for total CX3CR1 measurements only, PBS 1x was removed and replaced by permeabilizing solution (0.1% Triton X-100 in PBS 1x) for 10 min; for plasma membrane CX3CR1 measurements, samples were incubated with PBS 1x (without Triton X-100) for an equal amount of time. Once this incubation was over, intracellular peroxidases were quenched by incubation with 1% H_2_O_2_ (in PBS 1x) for 20 min. Quenching solution was removed and samples blocked with 1% BSA (in PBS 1x) for 1 h at room temperature. Blocking solution was discarded and cells incubated with anti-CX3CR1 (1:500; diluted in blocking solution) overnight, at 4 °C and the plate sealed with parafilm to prevent evaporation. In the following day, primary antibody solution was removed, samples washed thrice as described and incubated with HRP-conjugated anti-rabbit IgG (1:2000) for 2 h, at room temperature and protected from light exposure. Secondary antibody solution was removed, washed as described and samples incubated with the substrate solution (0.4 mg/mL OPD, 0.8 µL/mL in citrate buffer (111.2 mM NaH_2_PO_4_, 27 mM citric acid, pH 5.0)) for 30 min, at room temperature and protected from light. Chromogenic reaction was stopped by adding 5.5% H_2_SO_4_ solution to each well and absorbance measured after 5 min at 490 nm in the Multiskan GO plate reader (Thermo Fisher). For whole cell staining, the chromogenic solution was removed, cells washed five times with PBS 1x as described and incubated with 1% crystal violet solution for 30 min under gentle agitation. Crystal violet solution was removed, samples washed extensively with ddH_2_O to remove excessive amounts of dye, washed thrice with PBS 1x as described and incubated with 1% SDS solution for 1 h, under agitation, to solubilize intracellular crystal violet. Absorbance was read at 595 nm in the plate reader and this measurement was used to normalize CX3CR1 protein levels, according to the formula below:

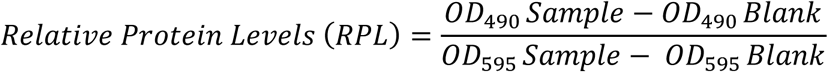

RPL was expressed relative to iMGs exposed to A_CM_ HCT (N.S.). Experiments were conducted twice with five replicates each.

### Mycoplasma testing

All cell lines were routinely tested for *Mycoplasma* contamination as described elsewhere (Molla Kazemiha et al., 2009) (universal *Mycoplasma* primers can be found in **Table S3**). Only *Mycoplasma*-free cells were used in this work.

### Statistical Analysis

Data were tested for Normal distribution using D’Agostino’s K-squared test. Except where otherwise specified, statistical analyses were carried out either by Two- or Three-way ANOVA, followed by Sidak’s multiple comparison test using the *GraphPad Prism* (v. 8.0.1) software. Synaptoneurosomes phagocytosis by iMGs were analyzed by multilevel mixed-effects linear regression with maximum likelihood estimation, followed by Sidak’s multiple comparison test in *R* (v. 4.3.1). Bioinformatics methods’ statistical analyses were conducted using each package’s built-in statistics in *R*. Unless otherwise stated, significance level was set at 0.05 (α = 0.05). Outliers were analyzed and removed using the ROUT Test (Q = 1%), when appropriate.

## RESULTS

### SCZ-sourced astrocytes display increased pro-inflammatory response following stimulation with TNF-α

Given the potential role for astrocytes in SCZ, we screened two transcriptomic (Akkouh et al., 2022; Szabo et al., 2021) and one proteomic (Trindade et al., 2023) studies that compared gene/protein expression changes between hiPSC-derived astrocytes from individuals with SCZ and neurotypical controls (HCT), in order to identify common biological processes associated with all three datasets. Interestingly, only the “cellular homeostasis” (GO:0019725) term overlapped among all three (**Figure S3A-B; Tables S4A-C**). Although no single candidate gene from the “cellular homeostasis” term overlapped among all three studies, three genes were present in at least two: CX3CL1 (the sole member of the CX3C chemokine family, also known as Fractalkine), NGFR (Nerve Growth Factor Receptor) and SLC24A2 (a calcium, potassium:sodium antiporter) (**Figures S3C-D**). CX3CL1 is associated with a wide range of biological processes that might be important in the context of SCZ, including microglial migration, immune response and synaptic pruning (**Figure S3E**), the latter having long been hypothesized as having a causative role in this disorder (Feinberg, 1982; Howes & Onwordi, 2023). Along with CX3CL1 (Paolicelli et al., 2011), classical complement components have also been implicated in synaptic elimination (Stevens et al., 2007; Yilmaz et al., 2021) (**Figure 1A**), while astrocyte-produced IL-33 has been shown to promote microglial-mediated synaptic uptake (Vainchtein et al., 2018). In addition, reactive astrocytes display increased expression of a wide range of cytokines with pleiotropic activities (Bernaus et al., 2020; Linnerbauer et al., 2020; Liu et al., 2011; Moraga, Spangler, Mendoza, & Garcia, 2014; Trindade et al., 2020), making them important targets for further investigation.

**Figure 1:**
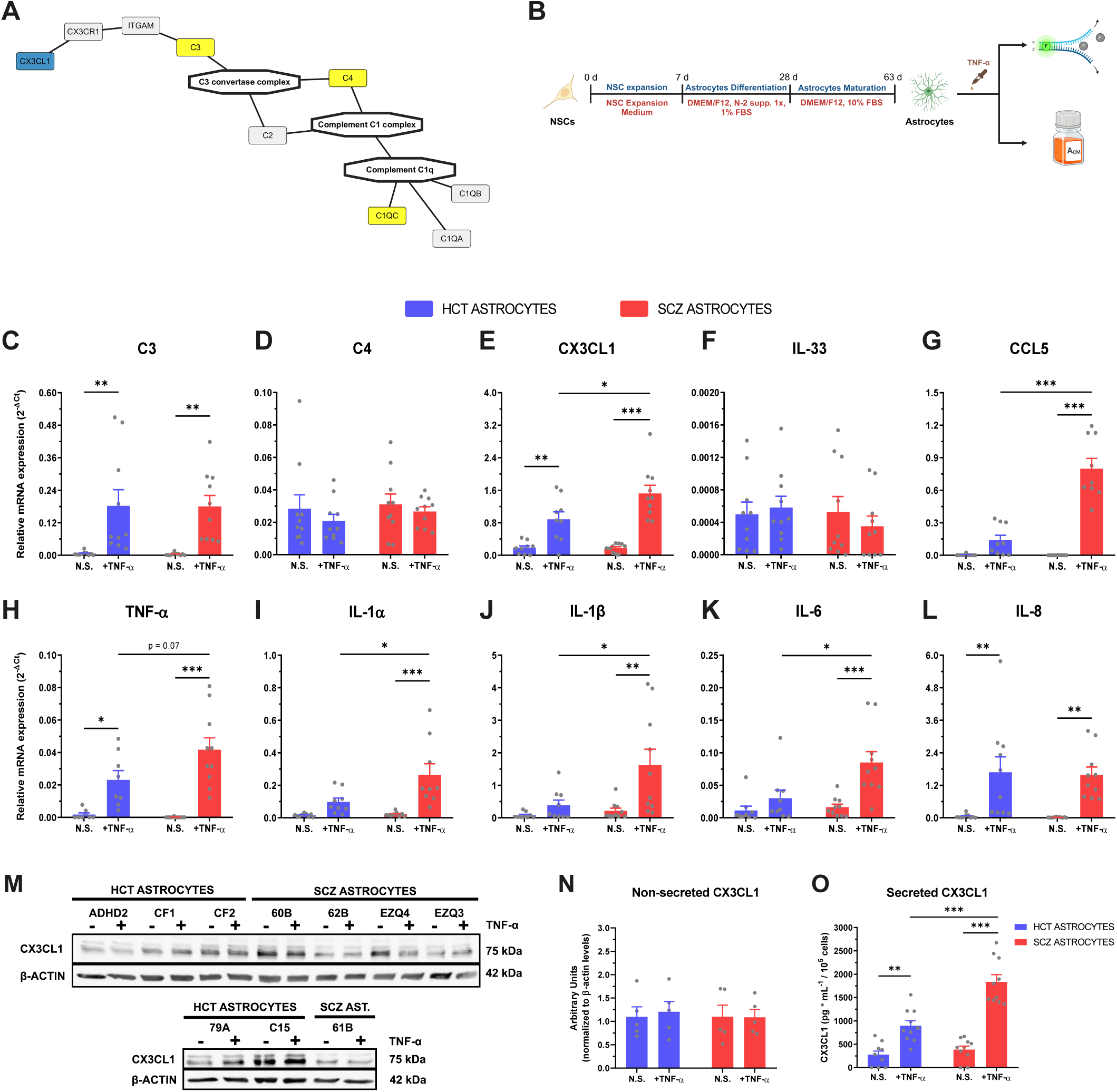
SCZ astrocytes showed stronger pro-inflammatory response upon TNF-α stimulation. **(A)** Synaptic pruning proteins network, highlighting CX3CL1 (blue) and classical complement components (yellow): C1q (subunit C), C3 and C4. Macromolecular complexes are drawn as hexagons and individual proteins as round rectangles. **(B)** Experimental design of hiPSC-derived astrocytes differentiation, stimulation with TNF-α (10 ng/mL) and downstream assays (RT-qPCR and conditioned media harvest for further applications). *Schematic picture was drawn using Biorender*. **(C-L)** mRNA expression of C3 **(C)**, C4 **(D),** CX3CL1 **(E),** IL-33 **(F),** CCL5 **(G),** TNF-α **(H),** IL-1α **(I),** IL-1β **(J)**, IL-6 **(K)** and IL-8 **(L)** in HCT and SCZ astrocytes stimulated with TNF-α. *n = 8-10 (two replicates x number of donor cell lines per group).* **(M-O)** Representative Western blot of non-secreted CX3CL1 in HCT and SCZ astrocytes stimulated with 10 ng/mL TNF-α **(M)** and its respective densitometric quantification **(N)**. *Representative blot out of three independent experiments*. *Data are expressed relative to non-stimulated HCT astrocytes. CX3CL1 levels were normalized relative to β-actin and displayed in arbitrary units. n = 5 (number of donor cell lines per group)*. **(O)** Secreted CX3CL1 levels in HCT and SCZ astrocytes stimulated with TNF-α. *n = 10 (two replicates x number of donor cell lines per group). Data were analyzed by Two-way ANOVA, followed by Sidak’s multiple comparison test. Bars represent Mean ± SEM. * p < 0.05, ** p < 0.01, *** p < 0.001*.

Taking in consideration the predicted involvement of inflammatory signaling in schizophrenia (Muller et al., 2015) and TNF-α ability in inducing astrocyte’s reactivity (Trindade et al., 2020), we decided to assay the expression of CX3CL1, IL-33, complement components and pro- and anti-inflammatory cytokines in hiPSC-derived astrocytes from five neurotypical (HCT) and five SCZ cases, upon stimulation with TNF-α (**Figure 1B**). Independent of cell donor diagnostic, CX3CL1 and C3 mRNAs were upregulated in astrocytes subjected to TNF-α stimulation, while C1q (subunit C) expression was not detected, and no change was observed in IL-33 and C4 expression (**Figure 1C-F**). Interestingly, CX3CL1 transcripts were higher in stimulated SCZ astrocytes compared to stimulated controls (**Figure 1E**). Moreover, TNF-α stimulation promoted the mRNA upregulation of the pro-inflammatory cytokines CCL5, TNF-α, IL-1α, IL-1β, IL-6 and IL-8, while the anti-inflammatory cytokines IL-4 and IL-10 were not detected (**Figure 1G-L**). Noteworthy, stimulated SCZ astrocytes displayed significantly augmented CCL5, IL-1α, IL-1β and IL-6 (and a trend towards increased TNF-α) mRNA levels compared to stimulated HCT astrocytes (**Figure 1G-K**).

Among these immune factors, CX3CL1 has been shown to be involved in a multitude of biological processes relevant in the context of SCZ **(Figure S3E)**. Thus, we sought to validate CX3CL1 mRNA expression findings **(Figure 1E)** at the protein level. TNF-α stimulation increased the production and secretion of CX3CL1 in both HCT and SCZ astrocytes, while no change in expression was observed at the non-secreted protein level **(Figure 1M-O)**. Remarkably, stimulated SCZ astrocytes showed an approximate two-fold augmentation in secreted CX3CL1 levels compared to stimulated control astrocytes **(Figure 1O).** Taken together, these data indicate that SCZ astrocytes responded stronger to TNF-α stimulation by producing greater levels of several pro-inflammatory cytokines and the chemokine CX3CL1, relative to HCT astrocytes.

### TNF-α-stimulated SCZ astrocytes promote a dysfunctional transcriptional response in microglial-like cells

Since many pro-inflammatory cytokines and CX3CL1 display pleiotropic functions, being capable of inducing several distinct responses in immune cells (Mecca, Giambanco, Donato, & Arcuri, 2018; Moraga et al., 2014) (**Figure S3E**), we queried potential biological processes that might be altered in microglial-like cells by reactive SCZ astrocytes. We generated iMGs from a neurotypical individual (**Figure S4**), incubated these cells with astrocyte conditioned media (A_CM_) harvested from HCT and SCZ astrocytes subjected to either vehicle or TNF-α stimulation (**Figure 1B**), and applied bulk RNA-seq to evaluate the transcriptional profile of these iMGs.

Principal component (PC) analysis showed that iMG samples exposed to A_CM_ SCZ (+TNF-α) clustered separately from iMGs cultured with any other A_CM_ in PC1; alternatively, PC2 segregated iMGs incubated with A_CM_ HCT (+TNF-α) from all other samples. Only the third principal component effectively split samples based on A_CM_ sourced from either HCT or SCZ astrocytes **(Figure 2A)**. Larger transcriptional differences were observed when iMGs were exposed to A_CM_ from TNF-α-stimulated astrocytes. Indeed, 414 differentially expressed genes (DEGs) (326 up- and 88 downregulated) were identified in iMGs + A_CM_ HCT (+TNF-α), while 847 DEGs (388 up- and 459 downregulated) were found in iMGs + A_CM_ SCZ (+TNF-α), and only 110 DEGs (83 up- and 27 downregulated) were detected in iMGs + A_CM_ SCZ (N.S.) **(Figure 2B-C; Tables S5A-C)**. Surprisingly, the iMGs + A_CM_ SCZ (N.S.) group shared more DEGs in common with iMGs + A_CM_ HCT (+TNF-α) (51 DEGs) than with iMGs + A_CM_ SCZ (+TNF-α) (14 DEGs) **(Figure 2B)**. These data corroborate the PC analysis results, since the first two groups are clustered closer in PC1 compared to the latter **(Figure 2A).**

**Figure 2:**
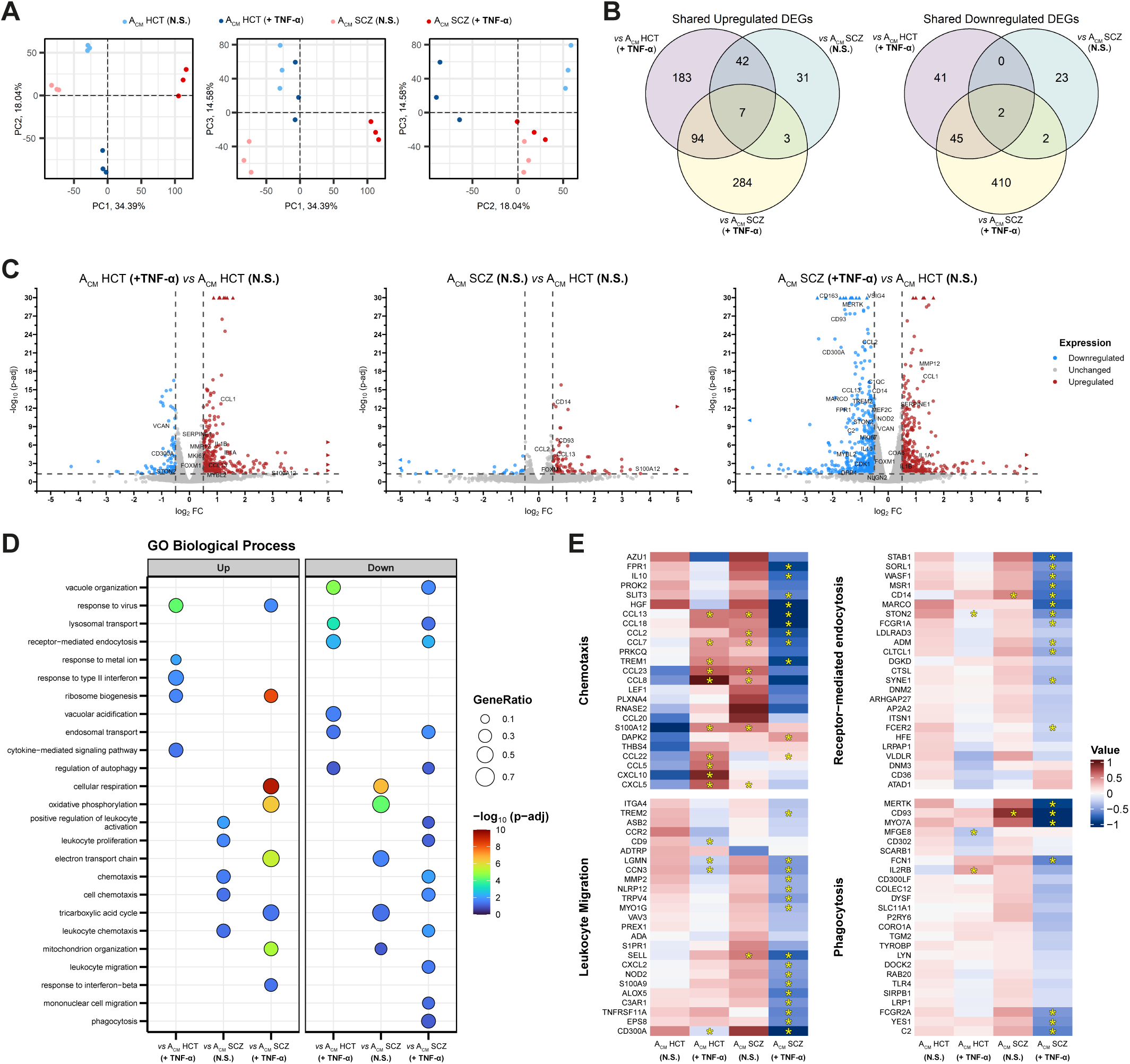
iMGs displayed a dysfunctional transcriptional profile after incubation with A_CM_ from TNF-α-stimulated SCZ astrocytes. **(A)** Principal component (PC) analysis of iMG samples incubated with all four distinct A_CM_ sources. (*left*) PC1 x PC2; (*middle*) PC1 x PC3; (*right*) PC2 x PC3. **(B)** Venn diagrams showing shared upregulated (*left*) and downregulated (*right*) DEGs between iMGs exposed to A_CM_ HCT (+TNF-α), A_CM_ SCZ (N.S.) or A_CM_ SCZ (+TNF-α). **(C)** Volcano plot depicting DEGs in iMGs cultured with: (*left*) A_CM_ HCT (+TNF-α) vs A_CM_ HCT (N.S.), (*middle*) A_CM_ SCZ (N.S.) vs A_CM_ HCT (N.S.), and (*right*) A_CM_ SCZ (+TNF-α) vs A_CM_ HCT (N.S.). Upregulated and downregulated DEGs are colored in red and blue, respectively. Circles and triangles indicate genes within and out of the plot axes range, respectively. **(D)** Gene-set enrichment analysis (GSEA) plot results presenting the most relevant GO Biological Processes associated with each experimental condition. *<u>vs</u> means that the indicated analysis is expressed relative to iMGs + A_CM_ HCT (N.S.).* **(E)** Heatmaps indicating relative gene expression levels in iMGs after incubation with A_CM_ HCT (N.S.), A_CM_ HCT (+TNF-α), A_CM_ SCZ (N.S.) or A_CM_ SCZ (+TNF-α). Each plot indicates the top 25 non-redundant genes associated with the following GO terms: *Leukocyte Migration, Chemotaxis, Phagocytosis* and *Receptor-mediated endocytosis*. Indicated genes may belong to more than one GO term. Data are displayed as mean-centered values*. Asterisks indicate differentially expressed genes (adjusted p-value < 0.05; |log_2_ Fold-change| ≥ 0.5)*.

Gene set enrichment analysis (GSEA) further explored the biological processes (GO terms) associated with each experimental condition. Incubation with A_CM_ SCZ (+TNF-α) led to the substantial downregulation of GO terms associated with several important microglial functions, including immune cell activation and proliferation, response to hypoxia, chemotaxis, cell migration and phagocytosis. A limited amount of downregulated GO terms was also shared with iMGs + A_CM_ HCT (+TNF-α), such as receptor-mediated endocytosis, regulation of autophagy, endosomal and lysosomal transport **(Figure 2D; Table S6A).** A closer inspection of the DEGs related to these downregulated GO terms unraveled that this molecular phenotype in iMGs + A_CM_ SCZ (+TNF-α) was mainly driven by decreased expression of cytokines and chemokines (e.g., *CCL2, CCL13, CCL18*), and immune and phagocytic receptors (e.g., *FPR1, TREM1, TREM2, MERTK, MARCO, IL2RA, IL2RB*) **(Figure 2E; Tables S5A-C and S6A).**

Conversely, both TNF-α-stimulated HCT and SCZ astrocytes induced the upregulation GO terms associated with ribosome biogenesis, response to virus and cellular response to interferon (IFN) in iMGs. Furthermore, TNF-α stimulation begot a shift in the direction of iMGs’ biological processes prompted by SCZ astrocytes. For example, while iMGs + A_CM_ SCZ (N.S.) displayed the upregulation of GO terms associated with chemotaxis, immune cell activation and proliferation, as well as the downregulation of processes related to mitochondrial metabolism, the contrary pattern was observed in iMGs + A_CM_ SCZ (+ TNF-α) **(Figure 2D, Table S6A)**.

To confirm these results, we carried out GSEA using KEGG pathways. Indeed, we confirmed that both A_CM_ HCT (+TNF-α) and A_CM_ SCZ (+TNF-α) induced the downregulation of lysosomal and autophagy pathways, while only the latter promoted a significant reduction in the endocytic pathway **(Figure S5; Table S6B)**. Noteworthy, TNF-α-stimulated SCZ astrocytes also triggered the upregulation of the p53 signaling pathway, reactive oxygen species production and multiple neurodegenerative disorders (e.g. Alzheimer’s and Parkinson’s Diseases) in iMGs, in marked contrast to iMGs + A_CM_ SCZ (N.S) **(Figure S5; Table S6B).**

To gain a better insight into how reactive SCZ astrocytes induce the downregulation of biological processes and pathways implicated in important microglial functions, we undertook a systems biology approach to establish whether their genes are organized into recognizable co-expression modules.

Weighted gene co-expression network analysis resolved 35 modules **(Figure 3A; Figure S6A-B, Table S7).** GO term overrepresentation analysis identified the biological processes associated with each module. The *brown* and *blueviolet* modules were largely enriched for terms involved in regulation of immune cell activation, autophagy, cell division and migration, chemotaxis, phagocytosis and endocytosis **(Figure 3B-C; Table S8).** Notably, these two modules were negatively correlated with iMGs incubated with A_CM_ SCZ (+TNF-α) and positively correlated with iMGs + A_CM_ SCZ (N.S.) **(Figure 3A).** Furthermore, the *antiquewhite2* and *lightgreen* modules, positively correlated with iMGs + A_CM_ SCZ (+TNF-α) samples, were enriched for GO terms linked to mitochondrial metabolism **(Figures 3A and S6C-D; Table S8).** On the other hand, the *darkolivegreen2* and *palevioletred* modules were positively correlated with iMGs cultured in the presence of A_CM_ HCT (+TNF-α) **(Figure 3A).** These two modules were mainly associated with biological processes related to response to cytokine and chemokine stimulation, as well as regulation of response to viruses and to IFN **(Figure 3D; Figure S6E; Table S8).** Together, these findings corroborate the GSEA results **(Figure 2D)** and suggest that SCZ genetic background and pro-inflammatory stimulation, such as by TNF-α, are simultaneously required in astrocytes to make microglial-like cells assume a dysfunctional molecular phenotype.

**Figure 3:**
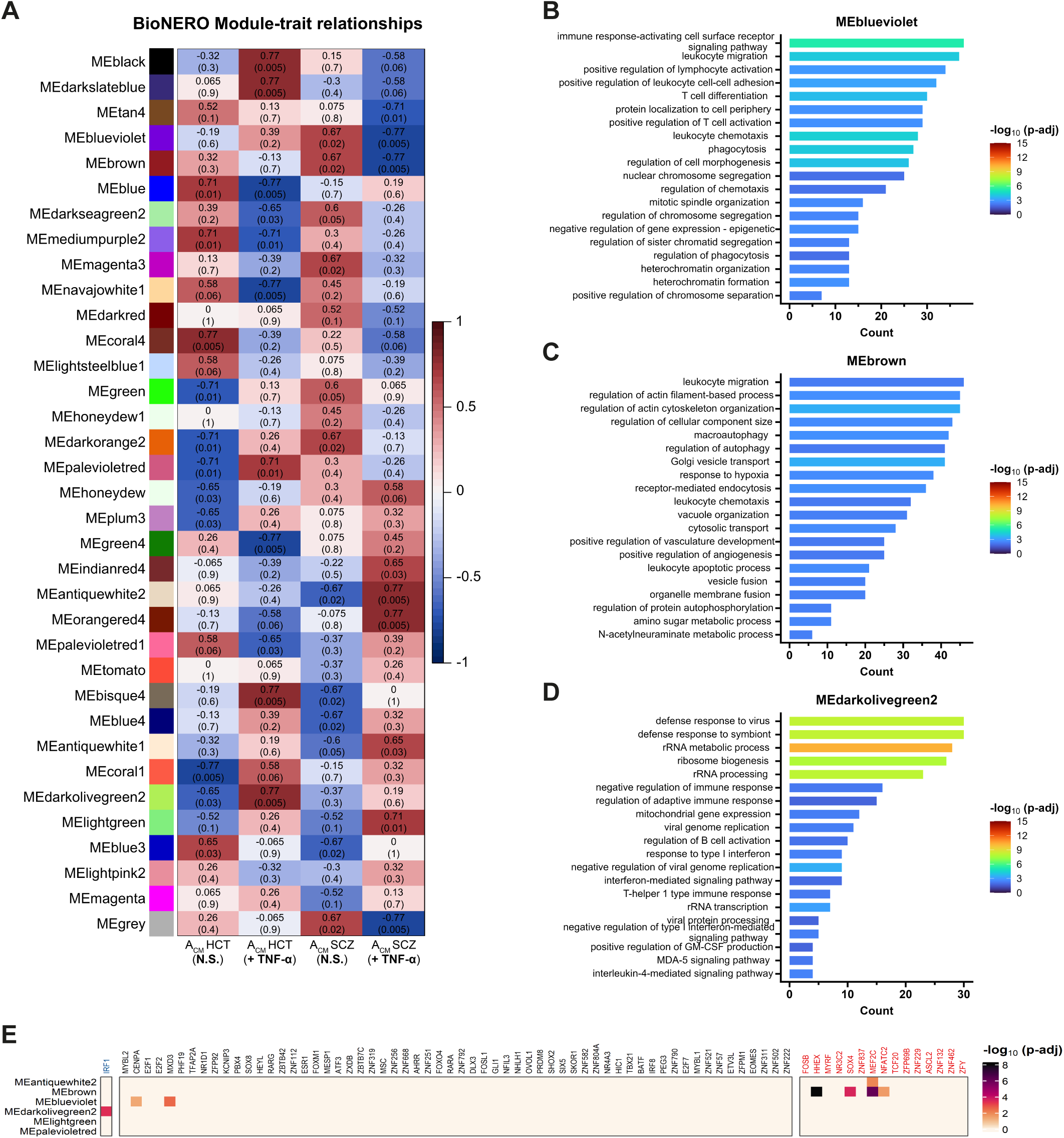
Identification of putative transcription factors that regulate co-expressed gene networks associated with impaired function in iMGs exposed to A_CM_ SCZ (+TNF-α). **(A)** Module-trait relationships heatmap displaying the correlation between WGCNA co-expression module eigengenes and each experimental condition (i.e., iMGs + indicated A_CM_). *Numbers in parentheses indicate adjusted p-values; above them are shown correlation values*. **(B-D)** Top 20 GO Biological Processes enriched in the *blueviolet* (B), *brown* (C) and *darkolivegreen2* (D) WGCNA modules. (E) *In silico* master regulator analysis depicting putative transcription factors (*columns*), whose predicted target genes showed significant overlap with the indicated WGCNA modules gene sets (*rows*). *Column names in blue or red indicate master regulators exclusively found in either iMGs + A_CM_ HCT (+TNF-α) or iMGs + A_CM_ SCZ (+TNF-α), respectively, while regulons shared by at least two experimental conditions are shown in black*.

### Master Regulator Analysis (MRA) reveals transcription factors involved in establishing the microglial dysfunctional response

Since co-expressed genes often share common regulators, including upstream transcription factors (TFs), we performed *in silico* master regulator analysis (MRA) using a comprehensive list of human TFs (Lambert et al., 2018) to identify prospective regulators in our dataset. A total of 161 regulons were identified, of which 126 were shared between at least two groups, 5 were exclusive to iMGs + A_CM_ HCT (+TNF-α) and 30 only found in iMGs + A_CM_ SCZ (+TNF-α) **(Figure S7A; Tables S9A-C).** Next, we calculated the predicted activity status of these transcription factors and only kept those whose activity were either considered as activated or repressed in at least one experimental group **(Figures S7B-D and S8; Tables S9A-C)**. To assess putative TFs regulating the co-expressed gene networks, we carefully cross-examined the target genes regulated by each TF and the gene sets pertaining to each WGCNA module. IRF1, whose activity status was predicted to be activated and was the only TF exclusive to iMGs + A_CM_ HCT (+TNF-α) **(Figures S7B and S8, Table S9A)**, was associated with the *darkolivegreen2* module **(Figure 3E)**, pointing to its possible role in inducing the expression of genes involved in response to cytokine, such as IFN, and chemokine stimulation in this experimental group. Interestingly, we found four (HHEX, SOX4, MEF2C and NFATC2) and two (CENPA and MXD3) TFs connected to the *brown* and *blueviolet* modules, respectively. MEF2C was also the only TF associated with the *antiquewhite2* module **(Figure 3E).** Except for SOX4, all these putative TFs had predicted repressed activity in the iMGs + A_CM_ SCZ (+TNF-α) group **(Figures S7D and S8, Table S9C)**. Among them, we observed that *MEF2C* mRNA expression was significantly reduced in iMGs cultured in the presence of A_CM_ SCZ (+TNF-α), while *NFATC2* and *CENPA* trended in the same direction **(Figure 2C and Table S5C)**. These data point to a putative role of MEF2C, NFATC2 and CENPA in promoting the dysfunctional microglial phenotype triggered by reactive SCZ astrocytes.

### Reactive SCZ astrocytes induce iMGs to assume a dystrophic/senescent-like phenotype

We were also puzzled by our results indicating that TNF-α-stimulated SCZ astrocytes promoted the upregulation of pathways associated with multiple neurodegenerative diseases, such as Alzheimer’s (AD) and Parkinson’s disease **(Figure S5 and Table S6B)**, and decided to investigate whether reactive SCZ astrocytes could prompt these dysfunctional microglia to assume a molecular profile akin to those observed in such neurological disorders. Hence, our dataset was compared to microglial transcriptional signatures from human prefrontal cortex (PFC) brain samples obtained from patients with AD (Prater et al., 2023). iMGs cultured with A_CM_ SCZ (N.S.) showed a positive association with the “interferon ELN (endolysosomal network)” signature, while iMG + A_CM_ HCT (+TNF-α) was associated with the “canonical inflammatory” and “cell cycle” ones. Remarkably, iMGs incubated with A_CM_ SCZ (+TNF-α) were found to be positively enriched for the “dystrophic” microglial phenotype **(Figure 4A)**.

**Figure 4:**
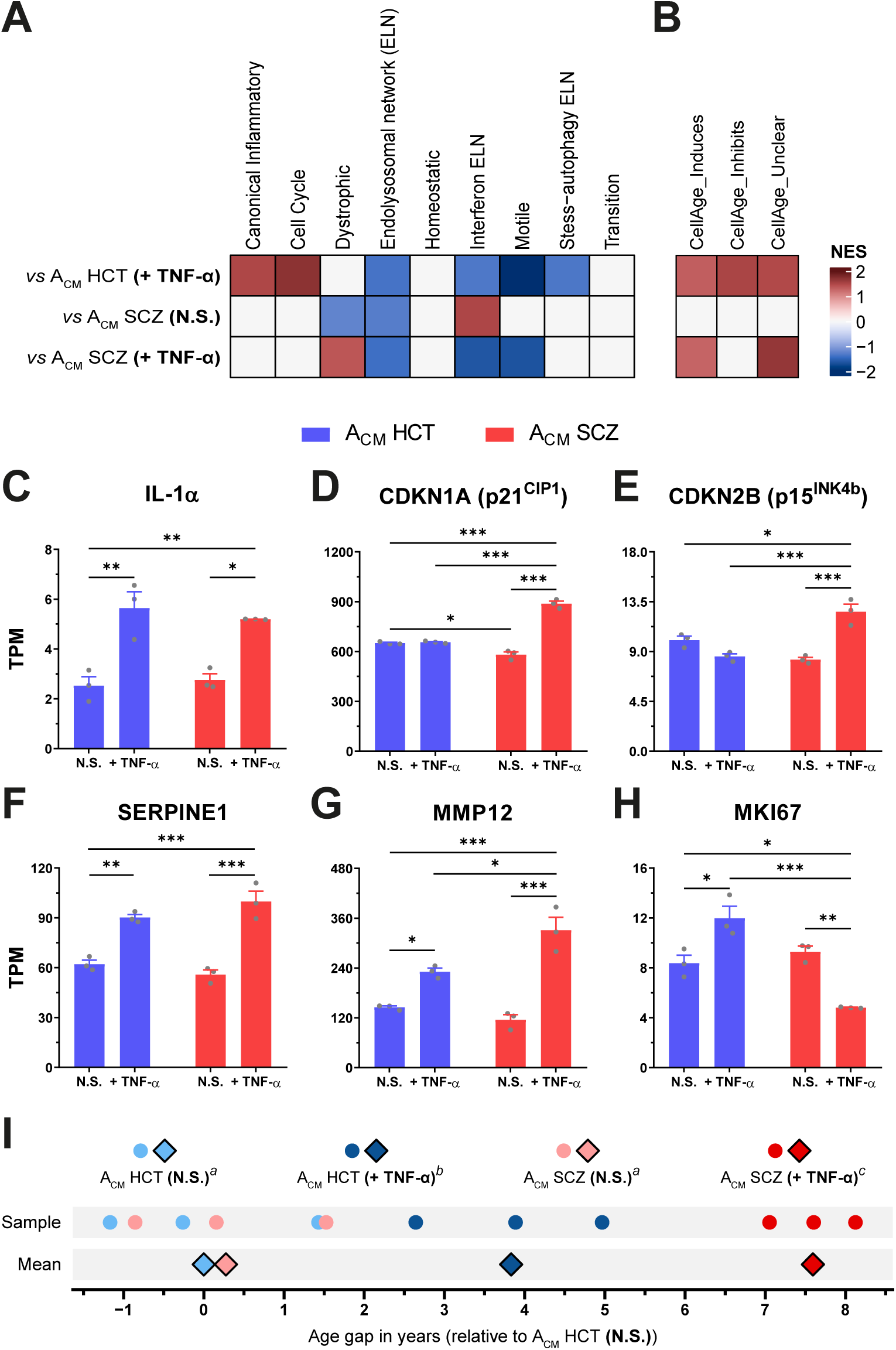
TNF-α-stimulated SCZ astrocytes promote a dystrophic microglial phenotype, leading to accelerated transcriptional age in iMGs. **(A)** iMGs’ molecular phenotypes identified after exposure to the indicated A_CM_ based on the human dorsolateral prefrontal cortex microglial transcriptional signatures described by Prater *et al* (2023) (Prater et al., 2023). **(B)** Microglial-like cells signature was assessed against the CellAge database (Avelar et al., 2020), whose genes are associated to either induction (*CellAge_Induces*), inhibition (*CellAge_Inhibits*) or have a context-dependent role (*CellAge_Unclear*) in promoting cellular senescence. *Only significant (p < 0.1) associations are colored in (A) and (B).* (C-H) mRNA expression of cellular senescence markers IL-1α **(C)**, CDKN1A **(D),** CDKN2B **(E),** MKI67 **(F),** MMP12 and SERPINE1 **(H)** in iMGs cultured with A_CM_ HCT (N.S.), A_CM_ HCT (+TNF-α), A_CM_ SCZ (N.S.) or A_CM_ SCZ (+TNF-α). *n = 3 (RNA-seq replicates). Data are shown in TPM (transcripts per million reads). Data were analyzed by Two-way ANOVA, followed by Sidak’s multiple comparison test. Bars represent Mean ± SEM. * p < 0.05, ** p < 0.01, *** p < 0.001.* (I) Transcriptional age gap calculated from the gene expression profile of iMGs incubated with the indicated A_CM_ and expressed relative to the mean of iMGs + A_CM_ HCT (N.S.) samples (mean was set as 0). *Data were analyzed by Two-way ANOVA, followed by Sidak’s multiple comparison test. Compact-letter display (*a, b *and* c*) shows the statistically significant groups (p < 0.05)*.

Dystrophic microglia display morphological, molecular and functional alterations characteristic of aged and senescent microglia (Angelova & Brown, 2019; Lopes, Sparks, & Streit, 2008; Rim, You, Nahm, & Kwon, 2024; Shahidehpour et al., 2021; Streit, Sammons, Kuhns, & Sparks, 2004). Then, we evaluated whether A_CM_ SCZ (+TNF-α) induced a senescent-like state in iMGs by assessing their transcriptional profile against the CellAge database (Avelar et al., 2020). Interestingly, iMG + A_CM_ SCZ (+TNF-α) displayed a positive association with genes that either induce or have a context-dependent regulatory activity (“unclear” category) towards cellular senescence, whilst iMG + A_CM_ HCT (+TNF-α) was also positively enriched for genes that display an inhibitory role **(Figure 4B).** To confirm these results, we analyzed the expression of some senescence markers (Suryadevara et al., 2024) in our RNA-seq dataset. Curiously, reactive SCZ astrocytes induced the mRNA expression of more senescent markers (*IL-1α, CDKN1A, CDKN2B, SERPINE1* and *MMP12*) in iMGs than their reactive HCT counterparts (*IL-1α, SERPINE1* and *MMP12*), with the former displaying an even greater expression of *MMP12* than the latter **(Figure 4C-G)**. In addition, iMG + A_CM_ HCT (+TNF-α) showed the augmented expression of *MKI67,* a well-known proliferation marker, while the contrary was observed in iMG + A_CM_ SCZ (+TNF-α) **(Figure 4H)**, indicating a potential downregulation in cell proliferation, a cellular senescence hallmark (Suryadevara et al., 2024).

Finally, iMGs’ transcriptional age was calculated to verify whether reactive SCZ astrocytes promoted accelerated aging, thus potentially leading to this dystrophic microglial phenotype. Intriguingly, exposure to both A_CM_ HCT (+TNF-α) and A_CM_ SCZ (+TNF-α) significantly increased the transcriptional age of iMGs, while no difference was observed between A_CM_ SCZ (N.S.) and A_CM_ HCT (N.S.) (+0.275 vs 0.000 years-old). Nonetheless, this transcriptional age enhancement was almost two-fold greater in iMGs incubated with A_CM_ SCZ (+TNF-α) than A_CM_ HCT (+TNF-α) (+7.593 vs. +3.831 years-old) **(Figure 4I).** Taken together, these results suggest that reactive SCZ astrocytes trigger accelerated aging in iMGs, prompting them to assume a dystrophic/senescent-like phenotype.

### Reactive SCZ astrocytes impair iMGs’ synaptic engulfment

Phagocytosis is amongst the key functions impaired in dystrophic microglia (Angelova & Brown, 2019), whose GO term was downregulated in iMGs after exposure to A_CM_ SCZ (+TNF-α) **(Figure 2D).** Given the strong association between synaptic uptake by microglial cells and SCZ development, we decided to assess whether reactive SCZ astrocytes could affect microglial engulfment of synaptic material **(Figure 5A)**. To establish this assay, we first isolated synaptoneurosomes from mature hiPSC-derived neurons **(Figure S9A-E),** fluorescently-labeled them and added to iMG cultures. As can be seen in **Figure S9F** and **Video S1**, iMGs were able to successfully phagocytose labeled synaptoneurosomes. Next, we tested whether A_CM_ from reactive SCZ astrocytes affected iMGs’ phagocytosis **(Figure 5A)**. Interestingly, incubation with A_CM_ SCZ (+TNF-α) was the only condition capable of diminishing synaptic engulfment by iMGs **(Figure 5B-E, Figure S10 and S11A).**

**Figure 5:**
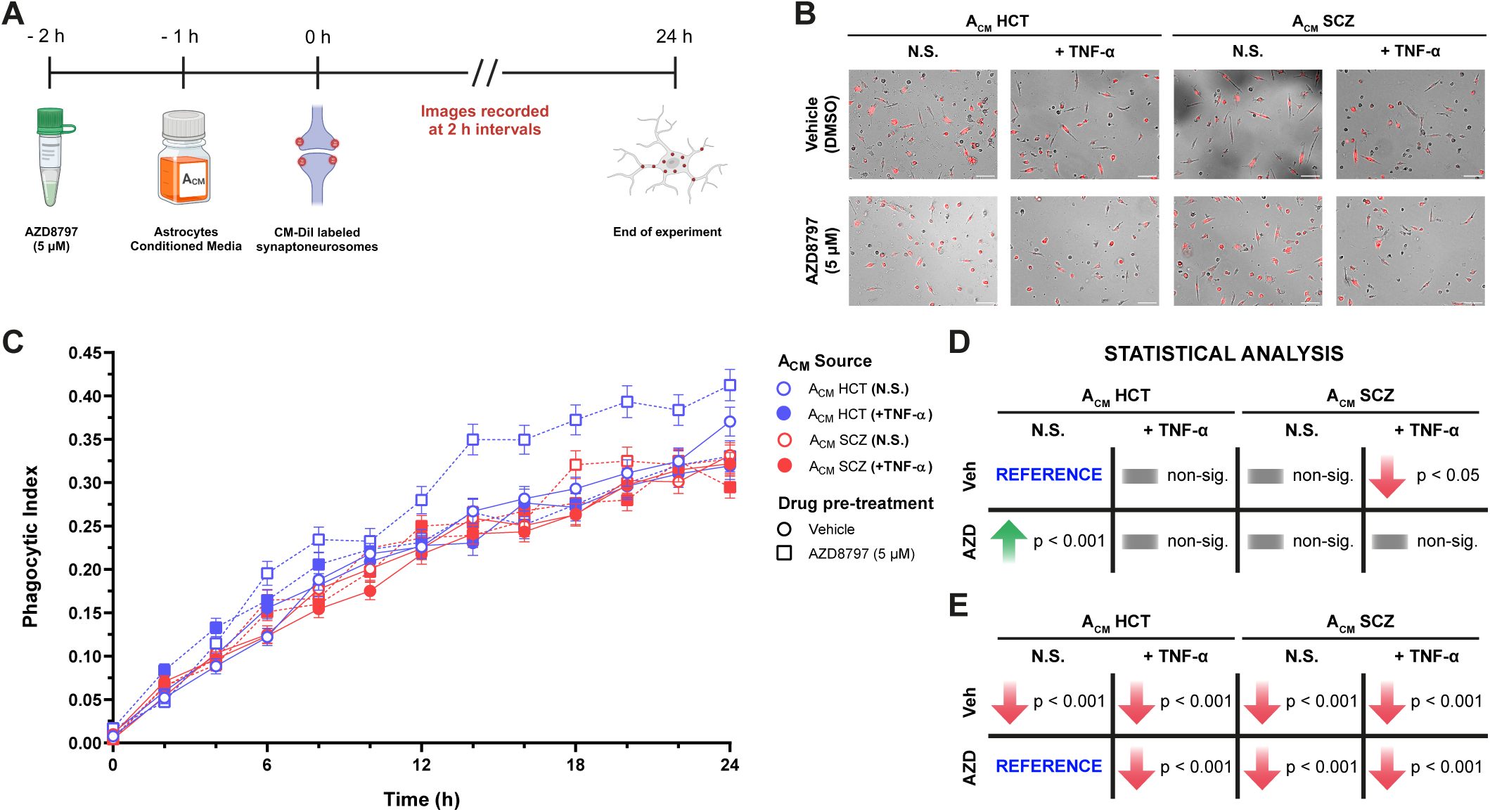
TNF-α-stimulated SCZ astrocytes impaired synaptoneurosomes engulfment by iMGs in a CX3CR1-independent manner. **(A)** Experimental design of synaptoneurosomes phagocytosis assay by iMGs after pre-treatment with AZD8797 (5 µM) and exposure to A_CM_. *Schematic picture was drawn on Biorender.* **(B)** iMGs (bright field) engulfing CM-DiI-labeled synaptoneurosomes (red) upon incubation with A_CM_ and after pre-treatment with the CX3CR1 antagonist AZD8797. Panel depicting 16 h time-point. *Scale bar = 100 µm.* **(C)** Quantification of synaptoneurosomes uptake by iMGs incubated with A_CM_ and pre-treated with AZD8797 depicted in **(B)** and **Figure S10**. Blue symbols: A_CM_ from HCT astrocytes; red symbols: A_CM_ from SCZ astrocytes; open symbols: A_CM_ from non-stimulated astrocytes; filled symbols: A_CM_ from TNF-α-stimulated astrocytes; circles and solid lines: vehicle (DMSO)-treated iMGs; squares and dashed lines: AZD8797-treated iMGs. *n = 355-744 (number of cells in 4 different fields of two independent experiments). Data were analyzed by Multilevel Mixed-effects linear regression, followed by Sidak’s multiple comparison test. Symbols represent Mean ± SEM. Pairwise significant comparisons are shown at the right side of the plot.* **(D-E)** Schematic representation depicting statistically significant differences depicting as reference groups vehicle-treated iMGs + ACM HCT (N.S.) **(D)** and AZD8797-treated iMGs + ACM HCT (N.S.) **(E)**, respectively.

Considering this result, we revisited our initial screening to select a potential target that could modulate this process **(Figure 1)**. We chose CX3CL1, as it has been previously shown to mediate synapse elimination (Paolicelli et al., 2011), and also displayed augmented secretion by stimulated SCZ astrocytes relative to stimulated controls **(Figure 1O)**. Then, to further evaluate its role, iMGs were incubated in the presence recombinant CX3CL1 (rCX3CL1), at approximately the same concentration secreted by reactive SCZ astrocytes **(Figure S11B).** Intriguingly, rCX3CL1 diminished synaptoneurosomes uptake by iMGs **(Figure S11C-D).** Next, we tested whether A_CM_ SCZ (+ TNF-α) required CX3CL1 signaling to lessen iMGs’ phagocytosis **(Figure 5A)**. Indeed, the pharmacological blockade of CX3CR1 by AZD8797 increased synaptoneurosomes engulfment relative to vehicle-treated iMGs under control conditions (A_CM_ HCT (N.S.)) **(Figure 5B-E, Figure S10, Figure S11E).** Nevertheless, CX3CR1 inhibition did not rescue the reduction in synaptoneurosomes phagocytosis by iMGs when incubated with A_CM_ SCZ (+TNF-α), nor did it lead to enhanced synaptic material uptake in iMGs cultured with either A_CM_ HCT (+TNF-α) or A_CM_ SCZ (N.S.), indicating that CX3CL1 was not responsible for promoting the observed phenotype **(Figure 5B-E)**. Overall, these data suggest that reactive SCZ astrocytes hinder microglial synaptic material phagocytosis. However, these effects on iMGs were CX3CL1-independent, not being reversed by the pharmacological inhibition of its receptor.

### TNF-α-stimulated SCZ astrocytes limit microglial migration

Cell migration is another phenomenon impaired in senescent microglia (Angelova & Brown, 2019), whose CX3CL1 involvement has already been demonstrated **(Figure S3E)**. Hence, we next queried whether reactive SCZ astrocytes affected microglial chemotaxis. Only iMGs exposure to A_CM_ from TNF-α-stimulated HCT astrocytes significantly enhanced microglial migration in a CX3CR1-dependent manner **(Figures 6A-B)**. Even though reactive SCZ astrocytes secreted almost twice as much CX3CL1 relative to their stimulated control counterparts, they failed to significantly induce microglial chemotaxis **(Figures 6A-B).**

**Figure 6:**
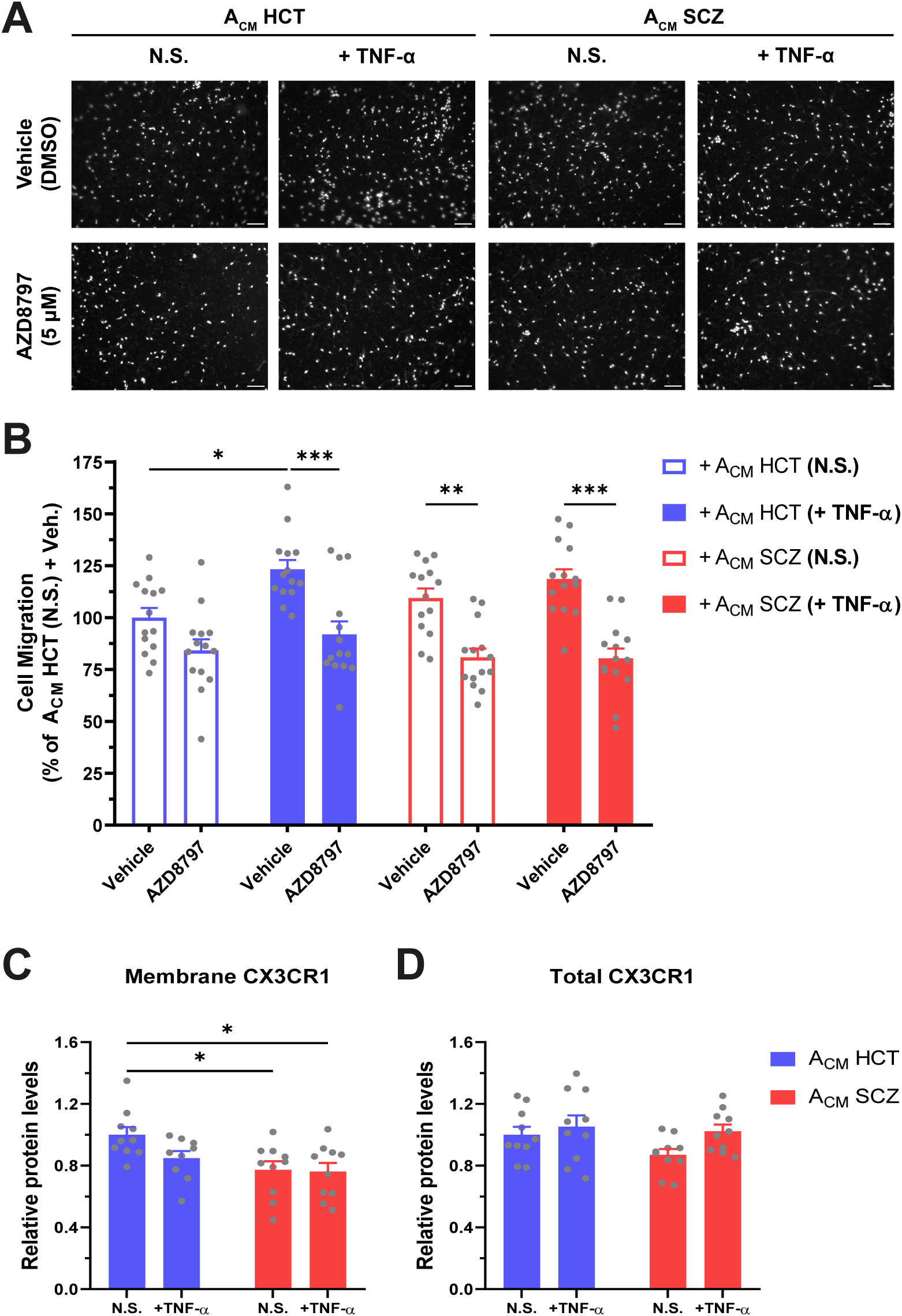
TNF-α-stimulated HCT and SCZ astrocytes promoted similar microglial migration in a CX3CR1-dependent manner. **(A)** Migrated iMGs after overnight incubation with A_CM_ and pre-treatment with either vehicle (DMSO) or the CX3CR1 antagonist AZD8797 (5 µM). Nuclei were stained with Hoechst (grayscale). *Scale bar = 100 µm.* **(B)** Quantification of iMGs’ migration shown in **(A)**. *n = 14 (4-5 fields per transwell x number of transwells per group). Data were normalized relative to the percentage of migrated iMGs in vehicle-treated cells cultured overnight in the presence of A_CM_ HCT (N.S.). Data were analyzed by Three-way ANOVA, followed by Sidak’s multiple comparison test. Bars represent Mean ± SEM.* **(C-D)** CX3CR1 relative protein levels measured in the plasma membrane **(C)** and in the whole cell **(D)** in iMGs incubated with A_CM_ for 24 h. *n = 9-10 (number of replicates in two independent experiments). Data were analyzed by Two-way ANOVA, followed by Sidak’s multiple comparison test. Bars represent Mean ± SEM. * p < 0.05, ** p < 0.01, *** p < 0.001*.

Intriguingly, an age-dependent decrease in CX3CR1 surface levels have been observed in human circulating monocytes (Seidler, Zimmermann, Bartneck, Trautwein, & Tacke, 2010). To assess whether this phenomenon could potentially limit iMGs’ response to reactive SCZ astrocytes-secreted CX3CL1, we measured the total and plasma membrane levels of CX3CR1 in iMGs exposed to A_CM_ for 24 h. SCZ A_CM_, but not A_CM_ HCT (+ TNF-α), diminished CX3CR1 plasma membrane content in iMGs, while total CX3CR1 levels remained unchanged **(Figure 6C-D)**. Taken together, these results indicate that the diminution of CX3CR1 plasma membrane levels prompted by reactive SCZ astrocytes act to restrict microglial-like cells migratory capabilities.

## DISCUSSION

Multiple lines of evidence link microglia and astrocytes to neuropathological mechanisms in SCZ (Gandal et al., 2018; Kim et al., 2024; Koskuvi et al., 2024; Szabo et al., 2021; Trindade et al., 2023; Uranova, Vikhreva, & Rakhmanova, 2021; Windrem et al., 2017), yet little is known about how these glial cells communicate and regulate each other’s function in this disorder. Here, we applied a combination of functional assays and transcriptomics to determine how SCZ astrocytes modulate key aspects of microglial biology. Reactive SCZ astrocytes displayed a stronger response to pro-inflammatory stimulation, that, in turn, prompted microglial-like cells to assume a dystrophic/senescent-like phenotype. Indeed, we observed that these dysfunctional microglia display features often found in aged microglia, including the upregulation of senescence markers (e.g., *CDKN1A, CDKN2B* and *MMP12*) and pathways (p53 signaling pathway), and the downregulation of several biological processes linked to cell proliferation, phagocytosis and chemotaxis. Our functional assays confirmed the occurrence of these last two phenomena by showing impairments in synaptic phagocytosis and limited microglial migration in a CX3CR1-independent and CX3CR1-dependent fashion, respectively.

Supporting this hypothesis, *post-mortem* findings reported augmented lipofuscin granules in microglia from younger and older individuals with SCZ, another characteristic cellular senescence feature (Uranova et al., 2021; Uranova, Vikhreva, Rakhmanova, & Orlovskaya, 2020). Furthermore, plenty of studies using anatomical, imaging, proteomic, metabolomic and epigenomic techniques have suggested that individuals with SCZ display accelerated brain aging (Campeau et al., 2022; Caspi et al., 2024; Constantinides et al., 2023; Kaufmann et al., 2019; Lin et al., 2021; Ling et al., 2024; Stone et al., 2022; Tian et al., 2023). In addition, aged and senescent microglia, as well as other myeloid cells, have been shown to display impaired phagocytosis and cell migration (Angelova & Brown, 2019; Caldeira et al., 2014; Hearps et al., 2012; Sharma, 2021; Spittau, 2017; Yanguas-Casas, Crespo-Castrillo, Arevalo, & Garcia-Segura, 2020).

A potential link between senescent immune cells and the CX3CL1/CX3CRI axis was provided by Seidler et al. (2010). Indeed, the authors found that circulating monocytes harvested from older donors show diminished CX3CR1 surface expression, similar to what we observed in iMGs exposed to SCZ A_CM_ (Seidler et al., 2010). Given that CX3CR1 mRNA is downregulated in the blood and brain of patients with SCZ (Chamera, Szuster-Gluszczak, & Basta-Kaim, 2021; Gandal et al., 2018; Snijders et al., 2021), and knowing that the CX3CR1^A55T^ loss of function variant was identified as a SCZ risk factor (Ishizuka et al., 2017), we suggest that activated astrocytes contribute to the CX3CL1/CX3CR1 axis dysfunction in SCZ. Yet, one seeming contradiction in our results is that reactive SCZ astrocytes secreted twice the amount of CX3CL1 compared to HCT counterparts, but did not induce a strong migratory response in iMGs. One possible explanation lies in the reduction in the CX3CR1 cell surface content in iMGs when incubated with ACM SCZ (+TNF-α), along with the possible existence of alternate pathways leading to the transcriptional dysregulation of cell migration, as indicated by our results. Taken together, these results suggest that activated astrocytes disturb microglial CX3CL1/CX3CR1 signaling in SCZ.

We also employed computational methods to find putative transcription factors capable of promoting the microglial phenotypic transition observed here. We identified three potential candidates (MEF2C, NFATC2 and CENPA), whose predicted repressed activities in iMGs incubated with A_CM_ SCZ (+TNF-α) may be responsible for downregulating the expression of genes associated with immune cell activation and proliferation, chemotaxis and phagocytosis. Intriguingly, MEF2C and NFATC2 have already been implicated in SCZ (Cosgrove et al., 2021; Mitchell et al., 2018; Shimamoto-Mitsuyama et al., 2021), while decreased expression of MEFC2 and CENPA are associated with increased aging in microglia and p53-dependent senescence induction in fibroblasts, respectively (Deczkowska et al., 2017; Maehara, Takahashi, & Saitoh, 2010). These data suggest a potential mechanism explaining, at least in part, how reactive SCZ astrocytes prompt iMGs to assume a senescent-like state. However, a thorough validation about how each TF affects microglia in the context of SCZ is beyond the scope of this work.

Despite the relevance of the data shown here, some limitations of this work remain. Firstly, while we have provided strong evidence that reactive SCZ astrocytes induce a dystrophic phenotype in microglial-like cells, the same set of experiments should be conducted using microglia obtained from patients with SCZ to have a more complete picture of the crosstalk between these two glial cell types in the context of this disorder. Secondly, it is not entirely clear what developmental time frame is modelled in this study (i.e., whether our findings recapitulate events that take place before or after symptom onset). Further research using more suitable developmental models, such as microglia containing organoids from individuals with SCZ, should be carried out to address this point.

In summary, our results demonstrated that reactive SCZ astrocytes led to substantial alterations in microglial biology, both at functional and transcriptional levels. These findings should be viewed as part of a grander change in the immune response landscape of microglia. Together, they strengthen the hypothesis that SCZ development requires the simultaneous occurrence of a susceptible genetic background and external environmental stimuli (Muller et al., 2015).

## ACKNOWLEDGEMENTS

We acknowledge the contributions on technical support of Mr. Jamil Silvano de Oliveira, Mr. Samuel Tadeu Rocha, Mr. Leandro Costa Nascimento, Dr. João Trindade Marques, all current and former members of the Laboratory of Neurobiochemistry (UFMG, Belo Horizonte, Brazil) and all medical and nursery personnel at the Borges da Costa Clinic (UFMG Clinic’s Hospital, Belo Horizonte, Brazil). We also thank Dr. João Paulo Pereira de Almeida, Dr. Álvaro Gil Araújo Ferreiro and Ms. Sophie Turetsky Cohen for their valuable discussions and insights regarding this work. This research project was funded by Conselho Nacional de Desenvolvimento Científico e Tecnológico (CNPq) and Fundação de Amparo à Pesquisa do Estado de Minas Gerais (FAPEMIG) grants. P.L.C. was recipient of CNPq and Fulbright scholarships during this work and is currently recipient of FAPEMIG fellowship; J.P.S.L. and N.C.S. are recipients of CNPq graduate scholarships.

## AUTHOR’S CONTRIBUTION

P.L.C. and F.M.R. conceived and designed the study. P.L.C. performed candidate genes screening, total RNA extraction, hiPSCs-derived neurons characterization, synaptoneurosomal characterization, iMGs synaptoneurosomes engulfment analysis pipeline development and quantification, western blots, ELISA, cell-based ELISA and iMGs migration assay. G.V. differentiated hiPSCs to NSCs. G.V. and J.C.P.M. characterized hiPSCs and NSCs. P.L.C., J.P.S.L. and P.T. differentiated hiPSC-derived astrocytes. P.T. characterized hiPSC-derived astrocytes. P.L.C. and J.P.S.L. performed astrocytes stimulation, RT-qPCR experiments, hiPSC-derived neuronal differentiation, iMGs differentiation and characterization. P.L.C. and J.S.F. performed synaptoneurosomes isolation and synaptoneurosomal phagocytosis assay. J.P.S.L. carried out iMGs migration assay blind quantification. B.F.C. performed PBMC’s donor psychiatric evaluation. R.N. supervised PBMC’s donor candidate recruitment and helped to draft ethics committee protocol. N.C.S. drew whole blood from neurotypical donor. P.L.C., N.C.S. and R.C.C. performed PBMCs isolation from whole blood. R.C.C. and J.S.F. helped to standardize ELISA and cell-based ELISA protocols. P.L.C., P.T. and L.C. established iMGs differentiation protocol. M.H.B. supervised iMGs differentiation protocol establishment. Y.M.H.T. performed RNA quality assessment in Bioanalyzer. P.L.C. and I.J.S.F. prepared cDNA libraries prior to RNA-sequencing. P.L.C. and C.Y.L. conducted bioinformatic analyses and carried out statistical analyses. K.J.B. provided platforms for bioinformatic analysis. K.J.B. and F.M.R. supervised bioinformatic analysis. S.K.R. supervised hiPSC-derived NSCs and astrocytes differentiations and donated NSCs and astrocytes cell lines, generated in his laboratory. P.L.C. and F.M.R. prepared the manuscript. L.B.V. provided research funding. F.M.R. supervised this study and obtained research grant funding. All researchers involved in this study read, made their contribution to the final version of this manuscript and agreed to publish this work.

## DATA AVAILABILITY

Raw and normalized gene-level count data, R markdown and CellProfiler pipeline files can be found in https://github.com/plcardozo/Cardozo-et-al---iMG-transcriptomics. Unprocessed sequencing files (.fastq) and other data may become available upon request.

