## Supplementary Figures for "hiPSC-derived astrocytes from individuals with schizophrenia induce a dystrophic phenotype in microglial-like cells"

##### FIGURE S1

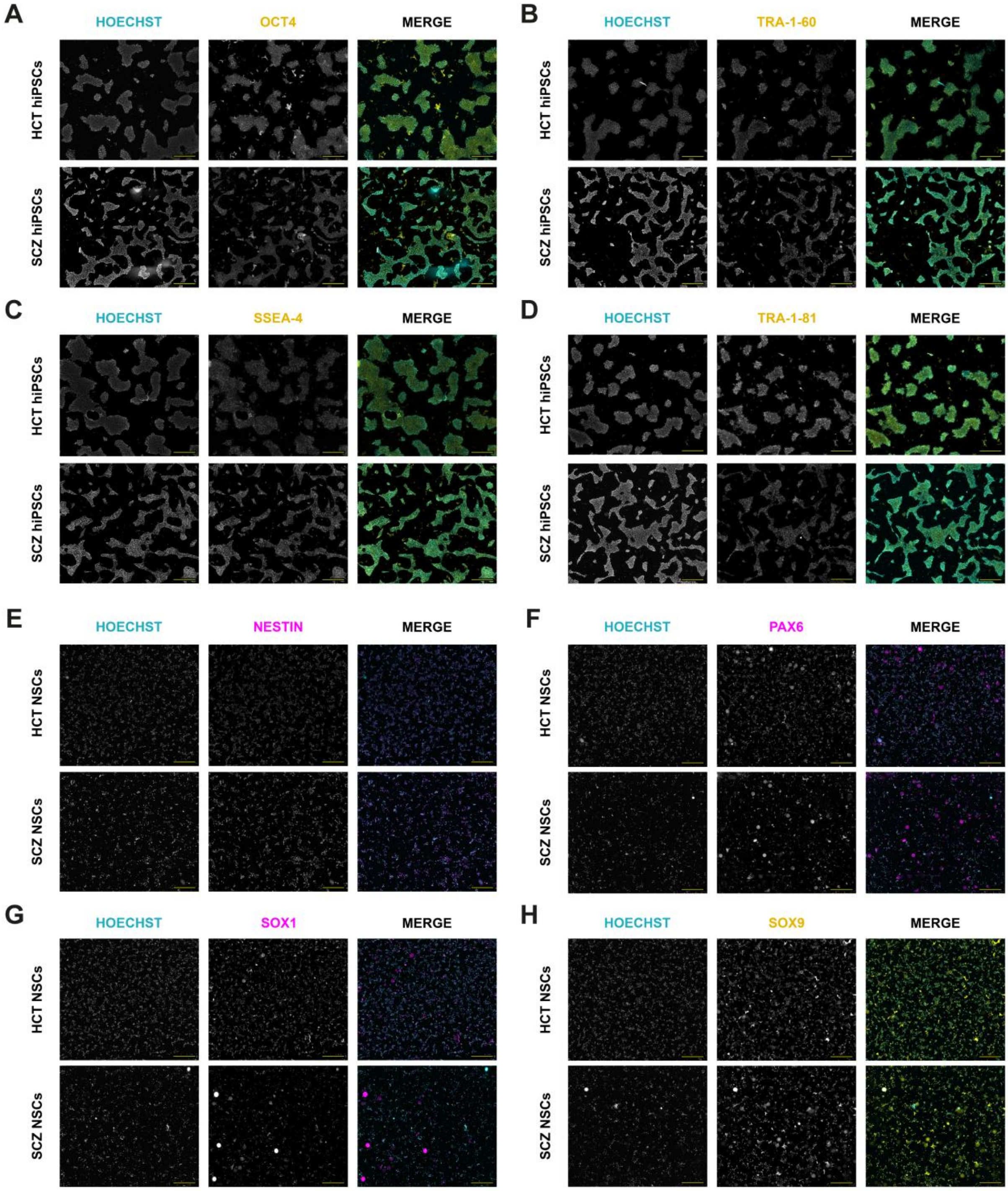


**FIGURE S1: hiPSCs and NSCs characterizations.** hiPSCs and hiPSC-derived NSCs sourced from HCT and SCZ individuals were characterized by immunostaining. **(A-D)** hiPSCs display positive staining for the pluripotency stem cell markers OCT4 (yellow; **A**), TRA-1-60 (yellow; **B**), SSEA-4 (yellow; **C**) and TRA-1-81 (yellow; **D**). **(E-H)** NSCs stains for the neural stem cell markers Nestin (magenta; **E**), PAX6 (magenta; **F**), SOX1 (magenta; **G**) and SOX9 (yellow; **H**). Nuclei were counterstained with Hoechst (cyan). *Scale bar = 700 µm.*

#### FIGURE S2

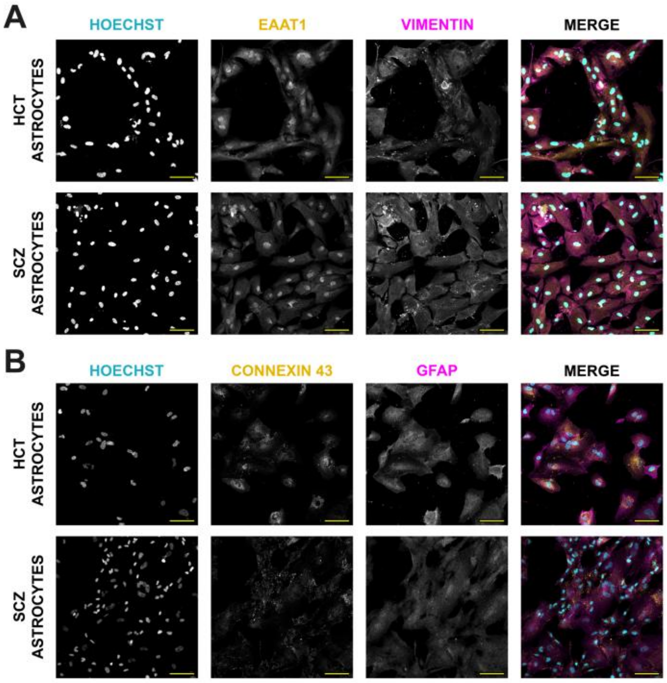


**FIGURE S2: hiPSC-derived astrocytes characterization. (A-B)** HCT and SCZ hiPSC-derived astrocytes were characterized by immunostaining. Astrocytes display positive staining for EAAT1 (yellow, **A**), Vimentin (magenta, **A**), Connexin 43 (yellow, **B**) and GFAP (magenta, **B**). Nuclei were counterstained with Hoechst (cyan). *Scale bar = 100 µm.*

#### FIGURE S3

A

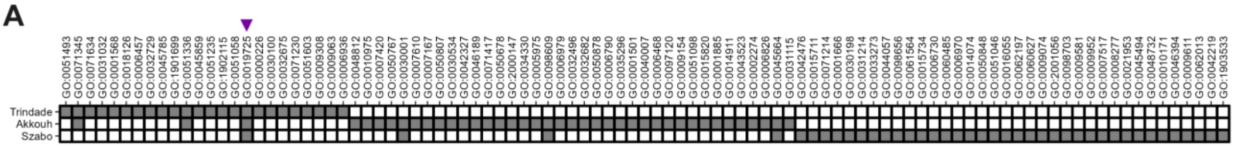


B

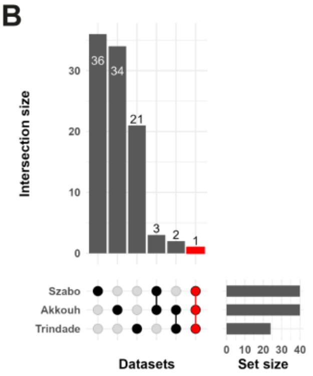


C

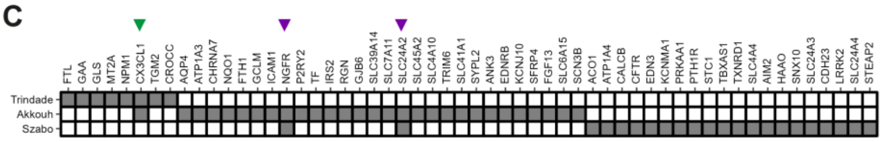


D

| Gene Symbol | Akkouh (Log₂ FC) | Szabo (Log₂ FC) | Trindade (Log₂ FC) | GO Molecular Function |
|---|---|---|---|---|
| CX3CL1 | ↑↑ (2.558) | - | ↑ (0.580) | Chemokine activity |
| NGFR | ↑↑ (2.143) | ↑↑ (3.341) | - | Nerve Growth Factor (NGF) binding |
| SLC24A2 | ↓↓ (-2.235) | ↑↑ (2.563) | - | Calcium, Potassium:Sodium antiporter activity |

E

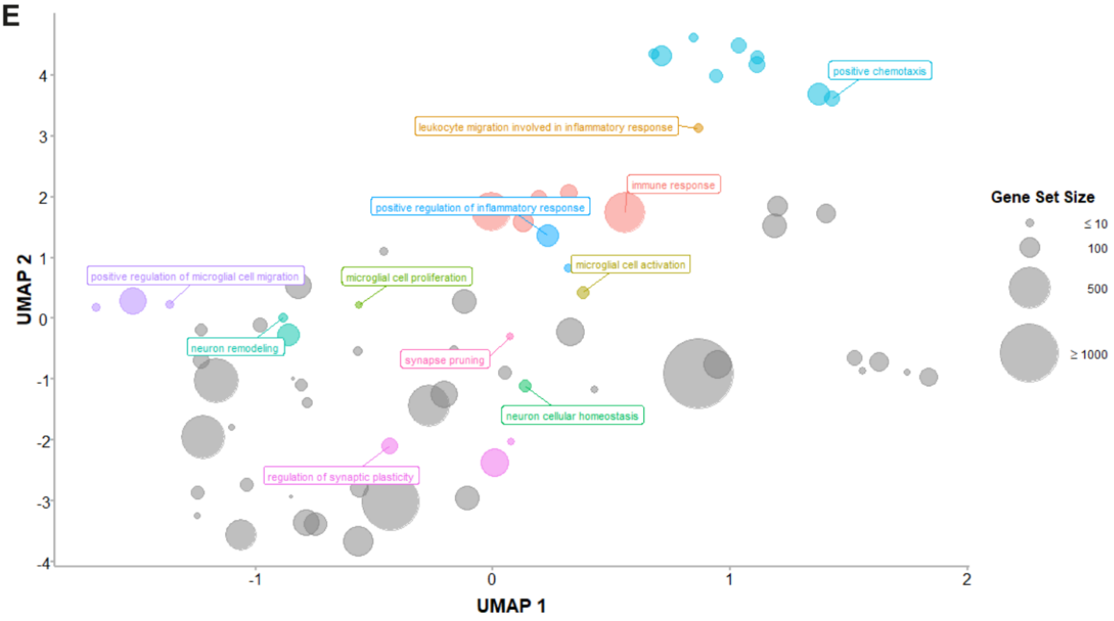


**FIGURE S3: Candidate gene screening suggested a potential role for astrocyte-produced CX3CL1 in schizophrenia. (A)** GO Biological Processes identified from Akkout *et al*, Szabo *et al* and Trindade *et al* datasets using *Metascape*. Filled and open squares indicate the enrichment or not of a given GO term in each dataset, respectively. Purple arrowhead points to GO:0019725 (cellular homeostasis), the only GO term to be enriched in the three studies. **(B)** Upset plot showing the number of overlapping enriched GO terms in all datasets. **(C)** Candidate genes extracted from GO:0019725 in each dataset. Purple arrowheads point to NGFR and SLC24A2 and green arrowhead points to CX3CL1, which has been chosen for subsequent investigation in the present study. **(D)** Table summarizing CX3CL1, NGFR, and SLC24A2 expression change in each dataset and their respective GO Molecular Function. **(E)** UMAP plot from semantic similarity analysis of all GO Biological Processes associated with CX3CL1. Circle size represents the gene set size of each depicted GO term. GO terms relevant to this study are colored and labeled accordingly.

#### FIGURE S4

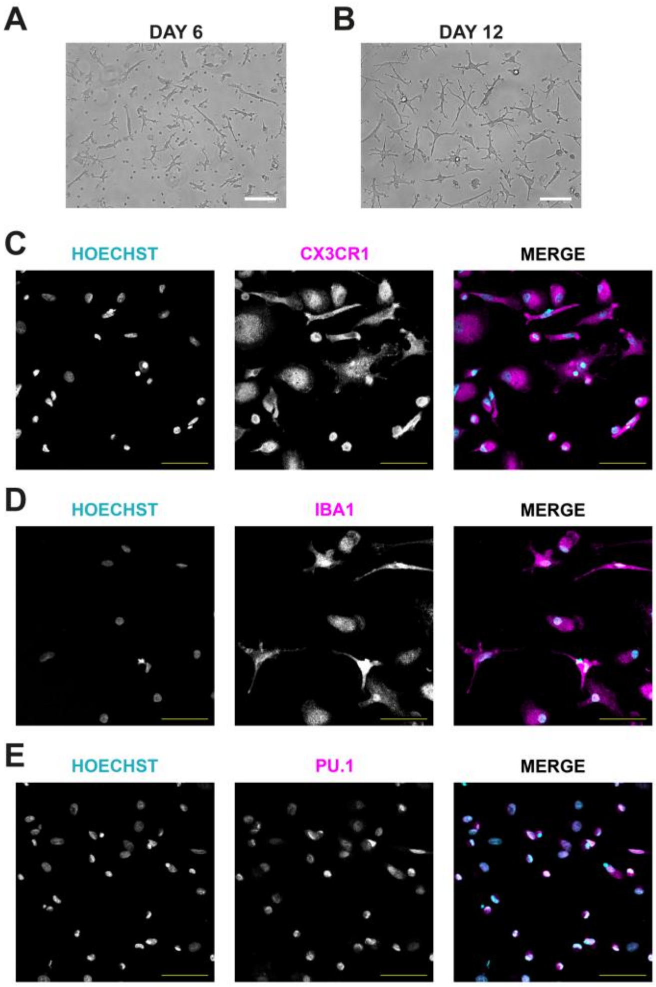


**FIGURE S4: Induced microglial-like cells (iMGs) characterization. (A-B)** Bright-field image of iMGs in their 6$^{th}$ **(A)** and 12$^{th}$ day of differentiation **(B)**, displaying a progressive ramified morphology. *Scale bar = 100 µm.* **(C-E)** 12-day differentiated iMGs were characterized by immunostaining. iMGs displayed positive staining for CX3CR1 (magenta, **C**), IBA1 (magenta, **D**) and PU.1 (magenta, **E**). Nuclei are counterstained with Hoechst (cyan). *Scale bar = 50 µm.*

**FIGURE S5**

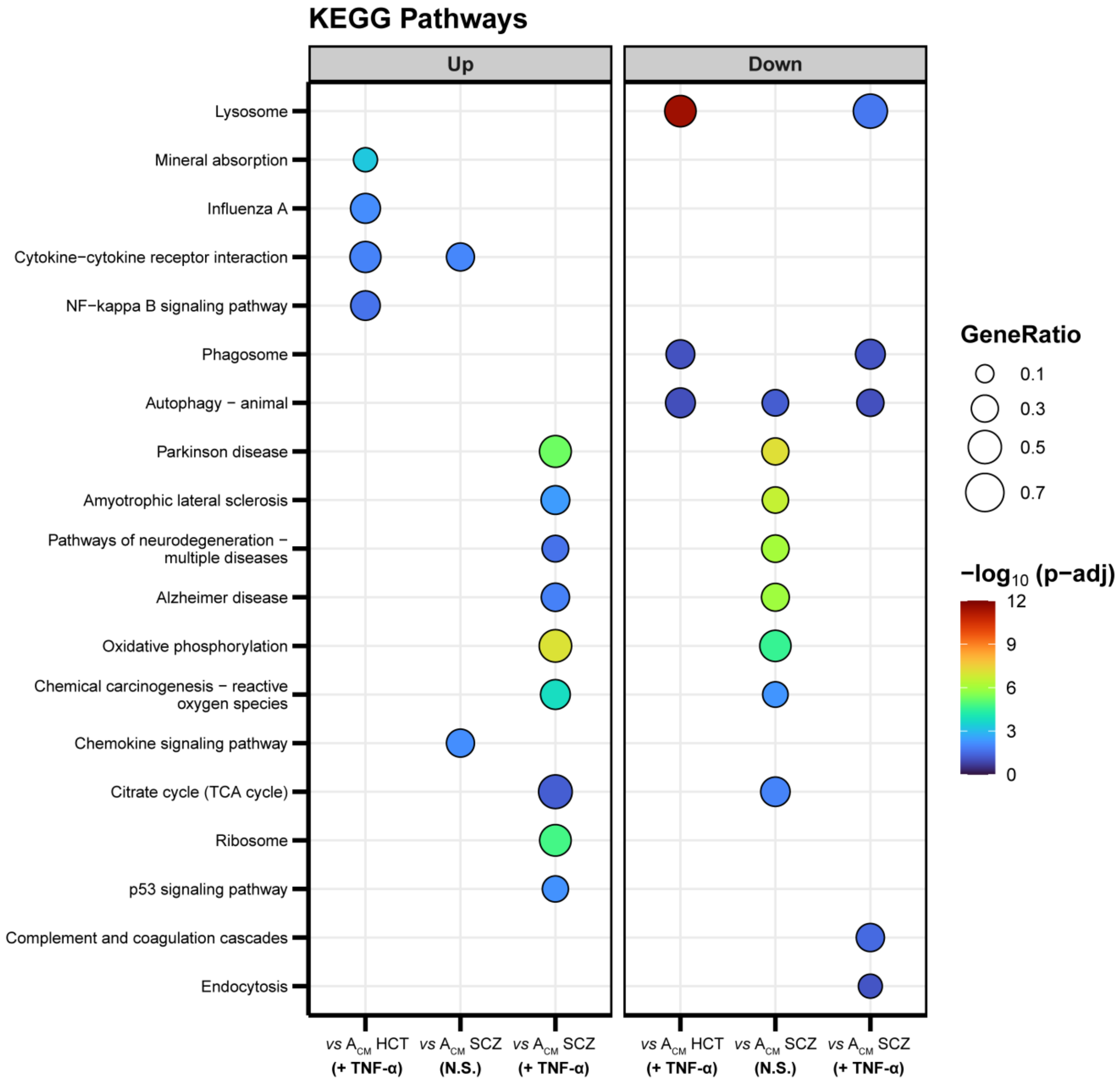


**FIGURE S5: TNF-α-stimulated SCZ astrocytes induced the upregulation of pathways related to neurodegenerative diseases in iMGs.** GSEA plot results showing the most relevant KEGG Pathways associated with each experimental condition. *vs means that the indicated analysis is expressed relative to iMGs +* $A_{CM}$ *HCT (N.S.).*

#### FIGURE S6

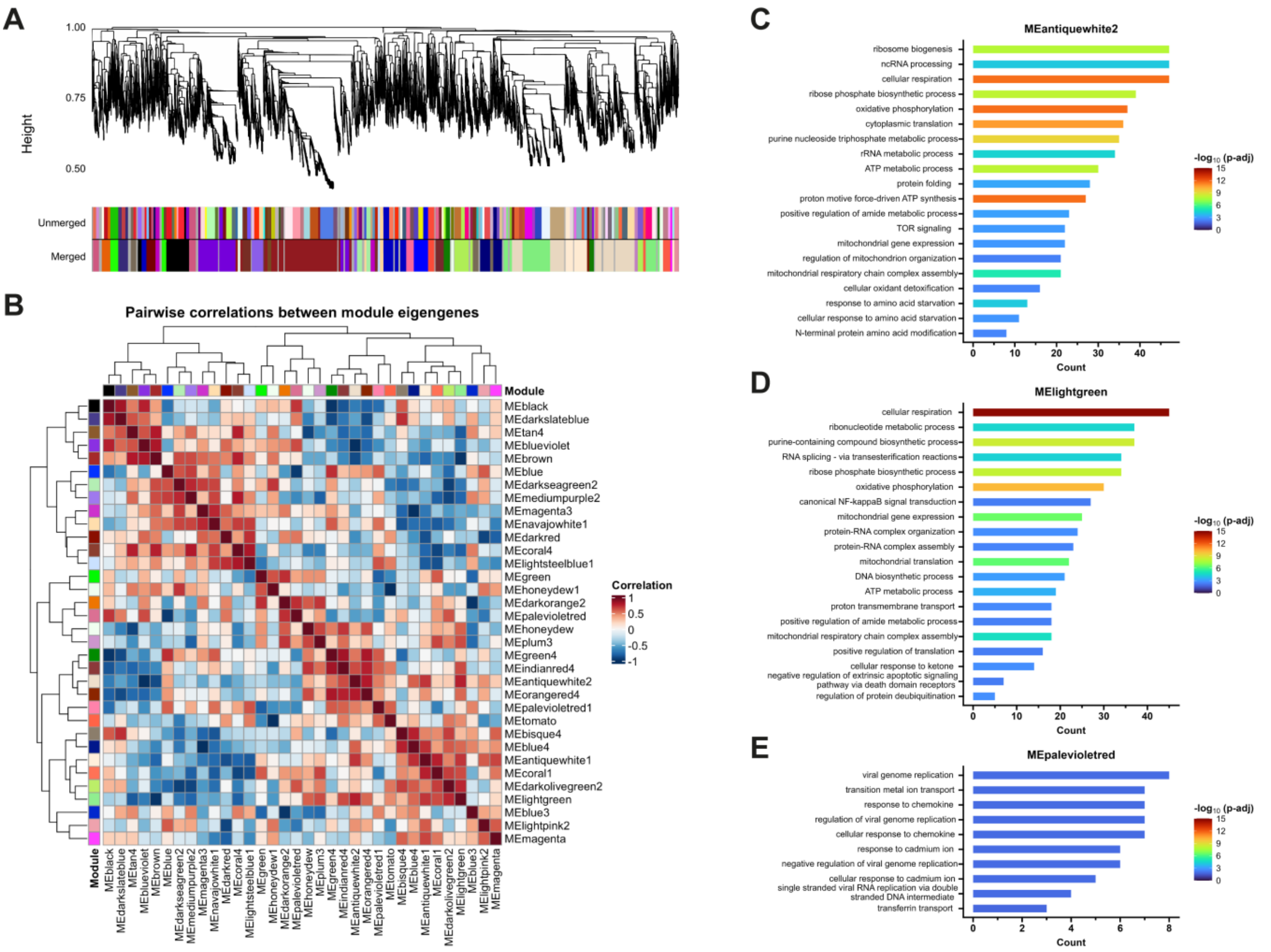


**FIGURE S6: WGCNA revealed modules enriched for GO terms involved in defense response against viruses and mitochondrial metabolism. (A)** Dendrogram depicting hierarchical clustering among identified co-expression modules. Modules with Spearman correlation greater than 0.8 were merged. **(B)** Heatmap showing the pairwise correlation between all 35 module eigengenes. **(C)** Top 20 GO Biological Processes enriched for the *antiquewhite2* module. **(D)** Top 20 GO Biological Processes enriched for the *lightgreen* module. **(E)** Top 10 GO Biological Processes enriched for the *palevioletred* module.

#### FIGURE S7

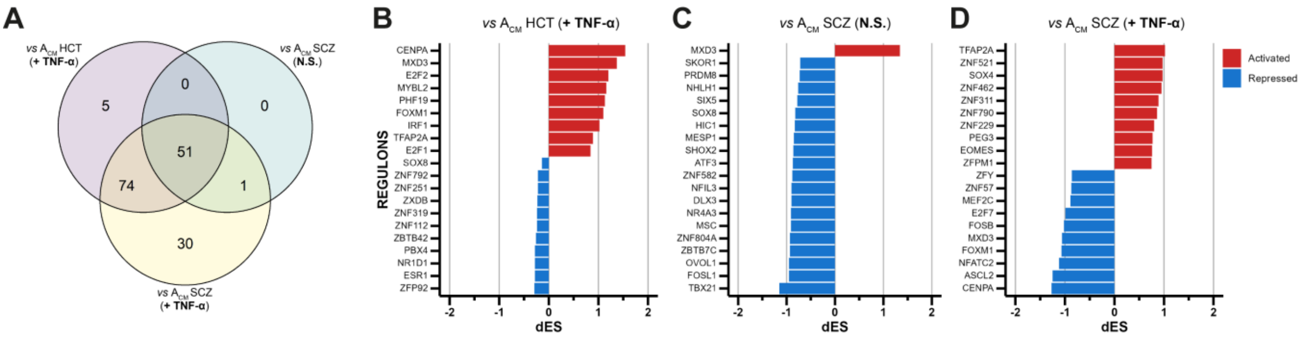


**FIGURE S7: Master Regulator Analysis identified putative transcription factors regulating the transcriptional response in iMGs incubated with $A_{CM}$. (A)** Venn diagram showing shared master regulators between iMGs exposed to $A_{CM}$ HCT (+TNF-α), $A_{CM}$ SCZ (N.S.), or $A_{CM}$ SCZ (+TNF-α). **(B-D)** Top and bottom 10 regulons ranked based on their activity status in iMGs + $A_{CM}$ HCT (+TNF-α) **(B)**, iMGs + $A_{CM}$ SCZ (N.S.) **(C),** and iMGs + $A_{CM}$ SCZ (+TNF-α) **(D)**. *dES: differential enrichment score.* *vs points that the indicated analysis is expressed relative to iMGs + $A_{CM}$ HCT (N.S.).*

### FIGURE S8

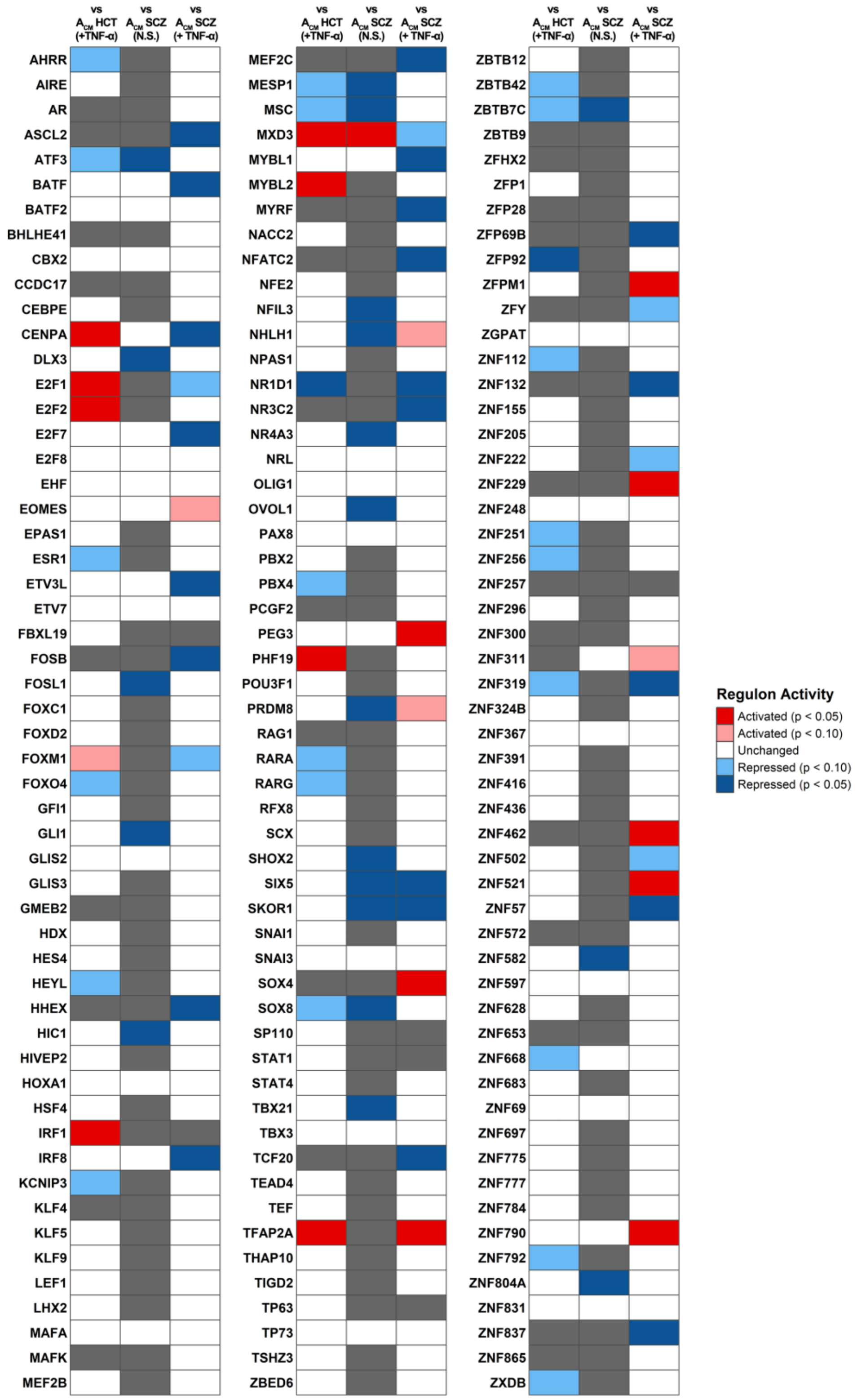


**FIGURE S8: Regulon activity of master regulators.** Regulon activity of master regulators is shown for each experimental group (iMGs incubated with $A_{CM}$ HCT (+TNF-α), $A_{CM}$ SCZ (N.S.) or $A_{CM}$ SCZ (+TNF-α)). *Light blue: repressed ($p < 0.1$); dark blue: repressed ($p < 0.05$); pale red: activated ($p < 0.1$); red: activated ($p < 0.05$); white: unchanged ($p \geq 0.1$); gray: regulon not significantly associated with that experimental group.* *<u>vs</u> points that the indicated analysis is expressed relative to iMGs + $A_{CM}$ HCT (N.S.).*

#### FIGURE S9

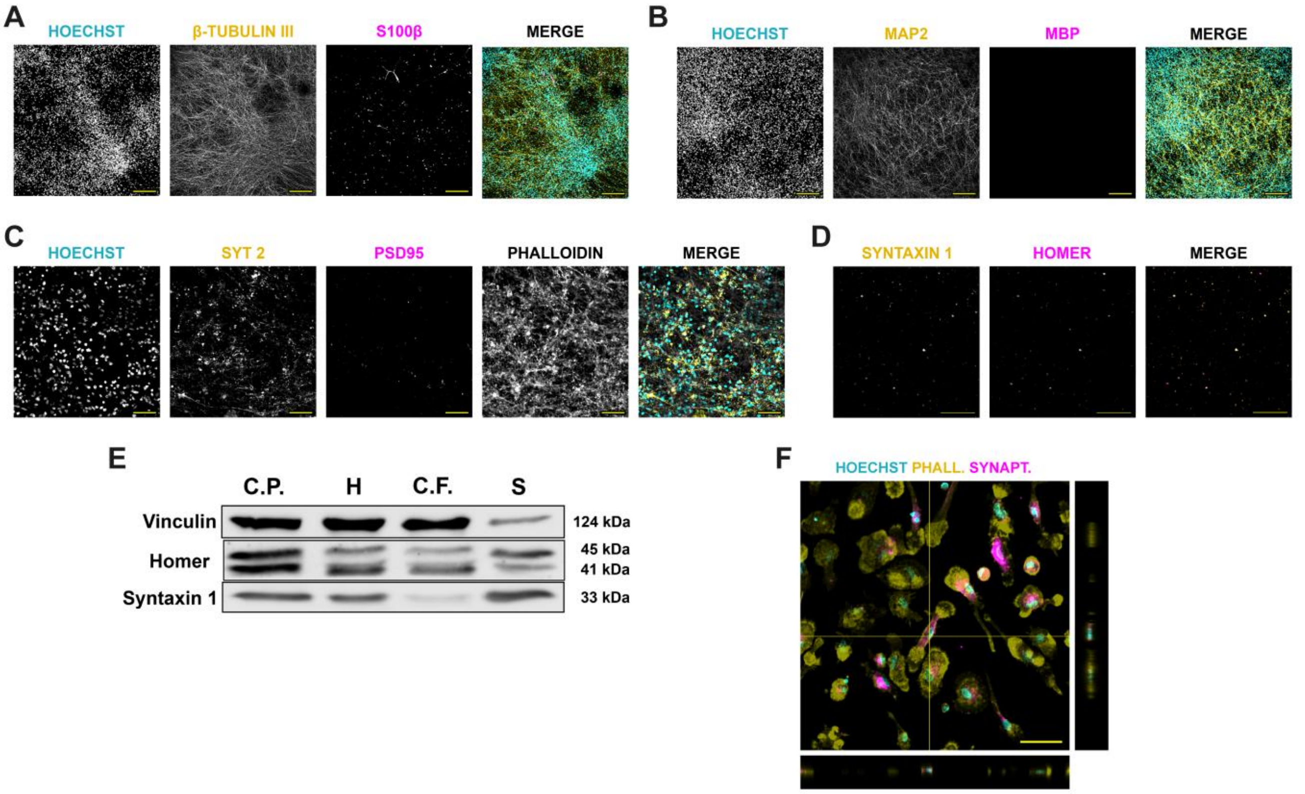


**FIGURE S9: Synaptoneurosomes isolation from hiPSC-derived neurons and their engulfment by iMGs. (A-B)** 60 days hiPSC-derived neuronal cultures stain for mature neuronal markers (β-tubulin III in **A**; MAP2 in **B**; yellow) and display few astrocytes (S100β in **A**; magenta) and no oligodendrocytes (MBP in **B**; magenta). Nuclei are counterstained with Hoechst (cyan). *Scale bar = 200 µm.* **(C)** Mature hiPSC-derived neurons show positive staining for presynaptic (SYT 2: Synaptotagmin 2; yellow) and postsynaptic (PSD95; magenta) markers. Cells were stained with Phalloidin (grey), and nuclei were counterstained with Hoechst (cyan). *Scale bar = 50 µm.* **(D)** Isolated synaptoneurosomes stained for the presynaptic marker Syntaxin 1 (yellow) and the postsynaptic marker Homer (magenta). *Scale bar = 50 µm.* **(E)** Western blot for Vinculin, Homer, and Syntaxin 1 of each fraction collected during synaptoneurosomes isolation procedure. *C.P.: cell debris pellet; H: homogenate; C.F.: cytosolic fraction; S: synaptoneurosomes.* **(F)** Representative image showing iMGs engulfing CM-DiI-labelled synaptoneurosomes (magenta), as shown in this orthogonal projection of a z-stack confocal microscopy image. iMGs are stained for Phalloidin (yellow) and nuclei are counterstained with Hoechst (cyan). *Scale bar = 50 µm.*

#### FIGURE S10

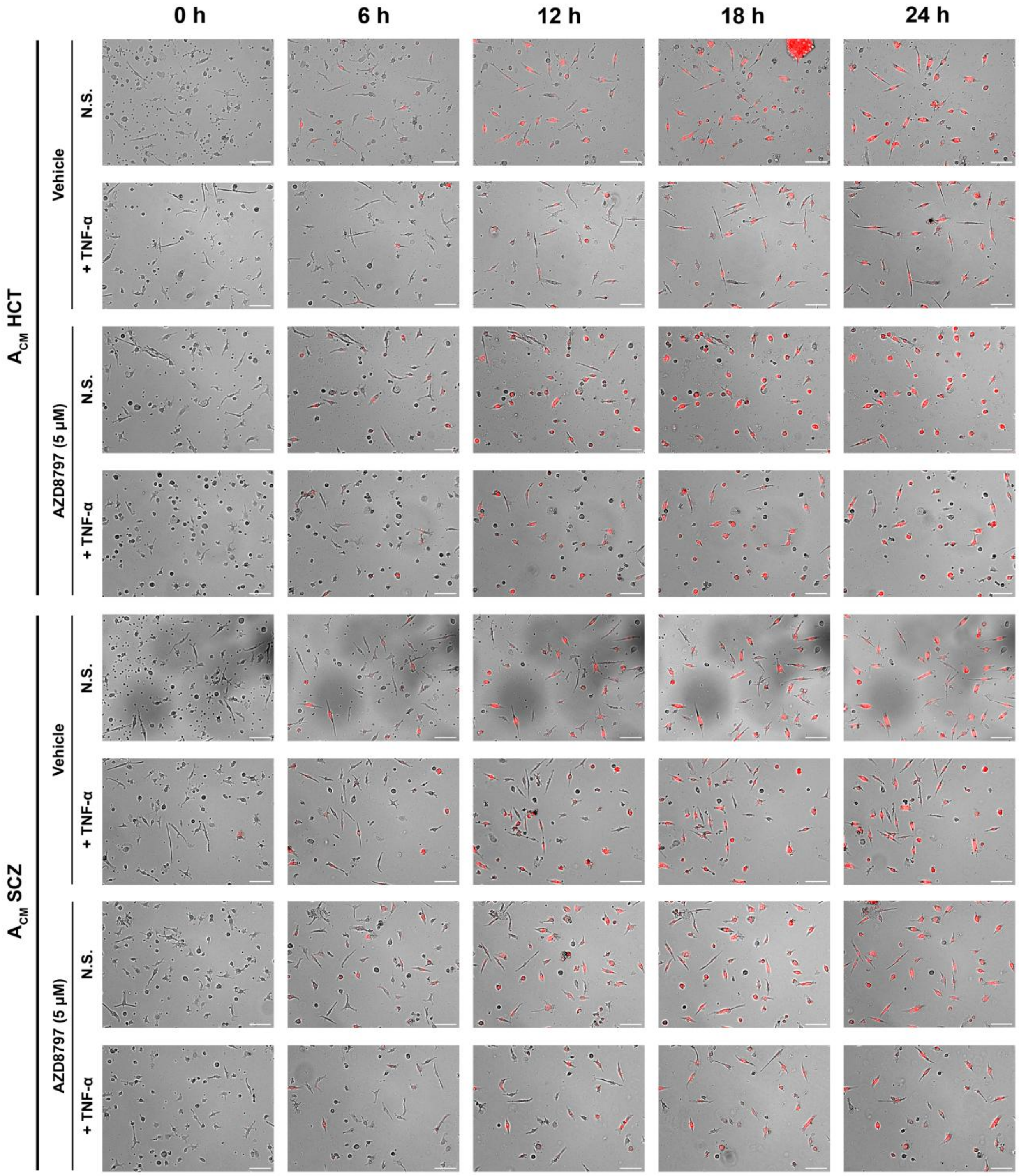


**FIGURE S10: iMGs engulfing fluorescently labelled synaptoneurosomes after pre-treatment with AZD8797 and following exposure to $A_{CM}$.** Full panel showing iMGs (bright field) phagocytosing CM-DiI-labelled synaptoneurosomes (red) upon incubation with $A_{CM}$ and pre-treatment with the CX3CR1 antagonist AZD8797 (5 µM). Panel depicting 0 h, 6 h, 12 h, 18 h, and 24 h time-points. *Scale bar = 100 µm.*

#### FIGURE S11

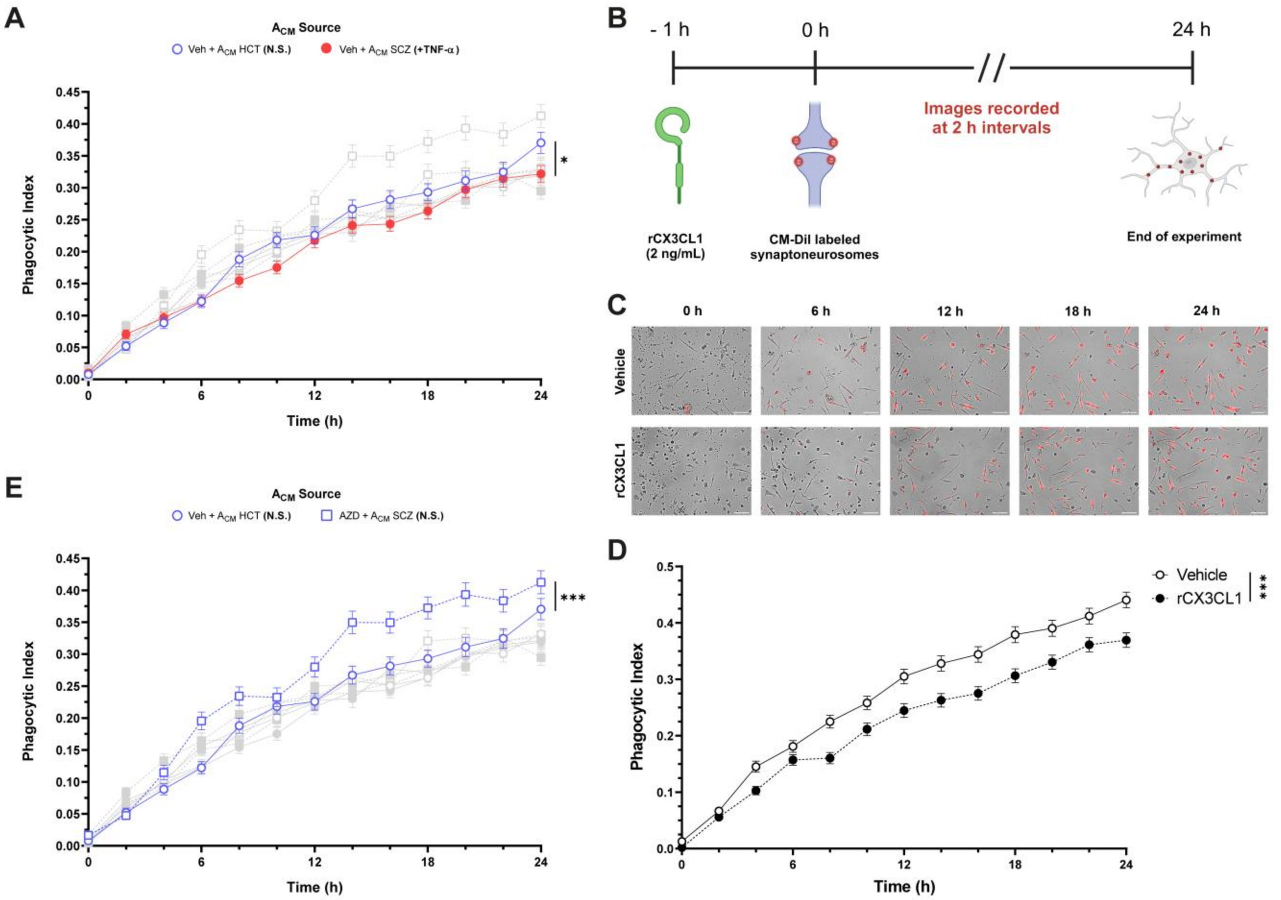


**FIGURE S11: CX3CL1 led to reduced synaptic engulfment by iMGs. (A)** Highlight of **Figure 5C** indicating the quantification of synaptoneurosomes engulfment in vehicle-treated iMGs incubated with either $A_{CM}$ HCT (N.S.) or $A_{CM}$ SCZ (+ TNF-α). **(B)** Experimental design of iMGs synaptoneurosomes phagocytosis assay upon exposure to recombinant CX3CL1 (2 ng/mL). *Schematic picture was drawn on Biorender.* **(C)** iMGs (bright field) engulfing CM-DiI-labeled synaptoneurosomes (red) during incubation with rCX3CL1. Panel depicting 0 h, 6 h, 12 h, 18 h and 24 h time-points. *Scale bar = 100 µm.* **(D)** Quantification of synaptoneurosomes phagocytosis by iMGs incubated with rCX3CL1 depicted in **(C).** Vehicle (BSA 0.1%; open circles); rCX3CL1 (2 ng/µL; filled circles). *n = 557-812 (number of cells in 4 different fields of two independent experiments).* **(E)** Highlight of **Figure 5C** indicating the quantification of synaptoneurosomes uptake in iMGs pre-treated with either vehicle or AZD8797 (5 µM) and culture with $A_{CM}$ HCT (N.S.). *Data were analyzed by Multilevel Mixed-effects linear regression, followed by Sidak's multiple comparison test. Symbols represent Mean ± SEM. $^*$ $p < 0.05$, $^{***}$ $p < 0.001$.*
